# Cytoplasmic nucleoporin foci are stress-sensitive, non-essential condensates in C. elegans

**DOI:** 10.1101/2022.08.22.504855

**Authors:** Laura Thomas, Basma Taleb Ismail, Peter Askjaer, Geraldine Seydoux

## Abstract

Nucleoporins (Nups) assemble nuclear pores that form the permeability barrier that separates nucleoplasm from cytoplasm. Nups have also been observed in cytoplasmic foci proposed to function as pore pre-assembly intermediates. Here we characterize the composition and incidence of cytoplasmic Nup foci in an intact animal, *C. elegans*. We find that, in young non-stressed animals, Nup foci only appear in developing sperm, oocytes, and embryos, tissues that express high Nup levels. The foci are condensates of highly cohesive FG-Nups that are maintained near their solubility limit in the cytoplasm by posttranslational modifications and chaperone activity. Only a minor fraction of FG-Nup molecules concentrate in Nup foci, which dissolve during M phase and are dispensable for nuclear pore assembly. Nup condensation is enhanced by stress and advancing age, and overexpression of a single FG-Nup in post-mitotic neurons is sufficient to induce ectopic condensation and organismal paralysis. Our results suggest that Nup foci are non-essential, “accidental”, and potentially toxic condensates whose assembly is actively suppressed in healthy cells.

## Introduction

In all eukaryotes, the double-membraned nuclear envelope partitions the nucleoplasm from the cytoplasm and material is exchanged between the two compartments by way of nuclear pore complexes. Pore complexes are composed of at least 30 distinct nucleoporins (Nups) arranged in biochemically stable subcomplexes (Figure 1A) (Cohen-Fix & Askjaer, 2017; Hampoelz *et al*, 2019a). Approximately two-thirds of Nups are essential to scaffold and anchor pore complexes to the nuclear envelope. The remaining one-third contain large phenylalanine/glycine (FG) rich domains that are highly intrinsically disordered. FG-Nups are enriched in the central channel of the pore and readily form multivalent interactions both *in vivo* and *in vitro* (Frey *et al*, 2006; Labokha *et al*, 2012; Patel *et al*, 2007; Xu & Powers, 2013). Cohesive interactions among FG-Nups are critical for the formation of the permeability barrier and FG-Nup hydrogels recapitulate nuclear pore selectivity *in vitro* (Frey & Görlich, 2007; Hülsmann *et al*, 2012; Ng *et al*, 2021; Schmidt & Görlich, 2015; Strawn *et al*, 2004). This has led to the “selective phase” model in which the permeability barrier is established by interactions among FG-Nups that form a phase separated network (Schmidt & Görlich, 2016; Ribbeck & Görlich, 2001).

**Figure 1.**
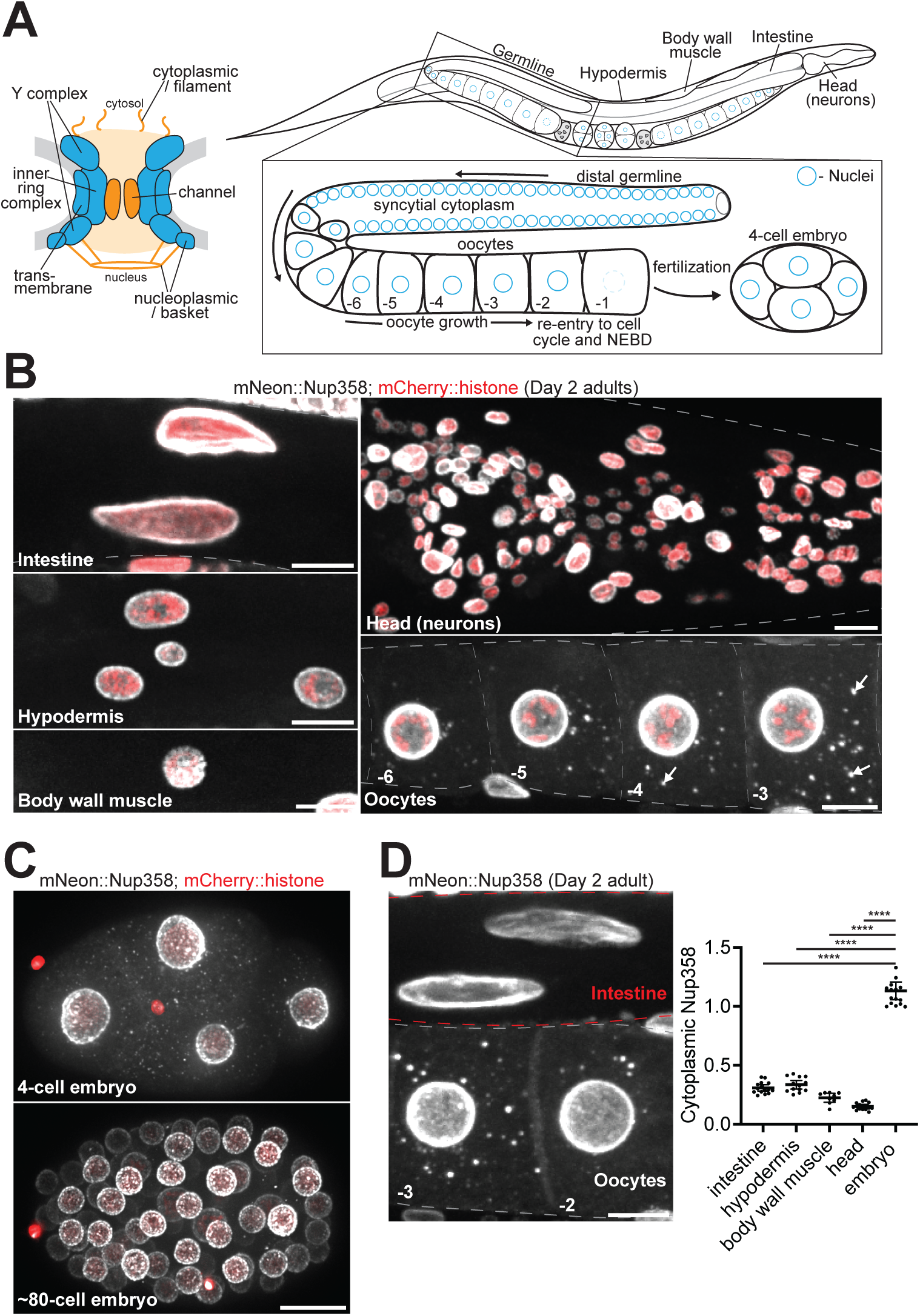
Cytoplasmic Nup foci are not present in somatic cells of young animals. A. Left: Schematic depicting the structure of a nuclear pore complex, which consists of ∼30 Nup proteins arranged in distinct subcomplexes. Blue subcomplexes are structural elements of the pore and include transmembrane Nups, the inner ring complex, and two copies of the Y complex. FG domain Nups are designated in orange and generate the permeability barrier of the central channel, and additionally localize to cytoplasmic filaments and the nuclear basket. Right: Schematic depicting the tissues and germline organization of a *C. elegans* adult hermaphrodite. Germ cell nuclei (designated in blue) proliferate in a syncytial cytoplasm before becoming enclosed by membrane to form individual oocytes. Oocytes arrest in meiosis I and grow in an assembly line-like fashion until induced by sperm signaling to re-enter the cell cycle in preparation for fertilization. B. Representative confocal micrographs of CRISPR-tagged mNeonGreen::Nup358 in the intestine, hypodermis, body wall muscle, head, and oocytes of Day 2 adult *C. elegans*. Nuclei are marked by a mCherry::histone transgene. White arrows denote cytoplasmic foci in oocytes. C. Representative confocal micrographs showing mNeonGreen::Nup358 in interphase 4-cell versus ∼80-cell embryos. D. Left: Representative confocal micrograph of mNeonGreen::Nup358 in -2 and -3 oocytes and intestinal cells of a Day 2 adult. Red dashed lines denote intestinal cells, gray dashed lines outline oocytes. Right: Quantification of cytoplasmic (soluble) mNeonGreen::Nup358 signal in intestinal, hypodermal, muscle, head (pharyngeal), or early (4-cell) embryonic cells as compared to that of the -1 oocyte. Values are normalized within the same animal so that the measurement for the -1 oocyte = 1.0. Error bars represent 95% CI for n > 7 animals. ****, P<0.0001. All images in this figure are maximum intensity projections. Scale bars = 10 μm.

In addition to their localization at the nuclear envelope, Nups have been observed in discrete cytoplasmic foci in oocytes, yeast cells, and in several other animal cell types cultured *in vitro* (Cordes *et al*, 1996; Raghunayakula *et al*, 2015; Ren *et al*, 2019; Colombi *et al*, 2013). Electron microscopy studies revealed that some cytoplasmic Nup foci correspond to annulate lamellae, a specialized subdomain of the endoplasmic reticulum (Kessel, 1989) proposed to function as a source of ready-made pore complexes in rapidly dividing cells (Hampoelz *et al*, 2016; Ren *et al*, 2019). The function of annulate lamellae, however, remains unclear and other studies have argued against a stockpiling function (Stafstrom & Staehelin, 1984; Onischenko *et al*, 2004). Cytoplasmic Nup foci have also been implicated in miRNA-mediated mRNA repression (Sahoo *et al*, 2017), nuclear pore inheritance (Colombi *et al*, 2013), and pore assembly by a condensate-based, non-canonical mechanism specific to oocytes (Hampoelz *et al*, 2019b).

Nups are also frequently enriched in pathological cytoplasmic inclusions that are hallmarks of neurodegenerative disease (Chandra & Lusk, 2022; Fallini *et al*, 2020; Hutten & Dormann, 2020), leading to the proposal that Nups become sequestered and depleted from nuclear pores under disease conditions (Gasset-Rosa *et al*, 2019; Zhang *et al*, 2018). Given the inherent propensity of FG-Nups to form multivalent networks, it is possible that Nup condensation may directly contribute to protein aggregation in disease. In support of this hypothesis, condensation of FG-Nup fusion oncogenes drives certain cancers (Chandra *et al*, 2022; Terlecki-Zaniewicz *et al*, 2021; Zhou & Yang, 2014), cytoplasmic Nup granules form upon loss of fragile X-related proteins (Agote-Aran *et al*, 2020), and cytoplasmic FG-Nups drive aggregation of TDP-43 in ALS/FTLD and following traumatic brain injury (Anderson *et al*, 2021; Gleixner *et al*, 2022). These observations suggest that Nups are not passive clients of cytoplasmic inclusions but rather active promoters of protein aggregation and disease progression.

Cytoplasmic Nup foci were reported previously in *C. elegans* oocytes and embryos (Pitt *et al*, 2000; Sheth *et al*, 2010; Patterson *et al*, 2011). Here we use the *C. elegans* model to systematically investigate the origin, regulation, and function of Nup foci. We show that Nup foci only contain FG-Nups and their direct interactors and form, in addition to oocytes and embryos, in developing sperm and in the somatic tissues of aged animals. In oocytes, the majority of FG-Nup molecules are maintained in a soluble cytoplasmic pool by posttranslational modifications and chaperone activity, with only a small minority (<3%) accumulating in foci. Condensation is enhanced by heat stress and FG-Nup overexpression, which when induced in neurons can disrupt nuclear pore assembly and lead to organismal paralysis. Together, our data suggest that Nup foci are non-essential, “accidental” byproducts of the natural tendency of FG-Nups to undergo condensation, which is essential to generate the permeability barrier of nuclear pores but must be suppressed in the cytoplasm to avoid premature and potentially toxic condensation.

## Results

### In young animals cytoplasmic Nup foci only assemble in growing oocytes, developing sperm, and early embryos

Nup358 and Nup88 have been reported in cytoplasmic Nup foci in oocytes, yeast, and a wide range of cultured cell types from different organisms (Sahoo *et al*, 2017; Hampoelz *et al*, 2019b; Colombi *et al*, 2013; Raghunayakula *et al*, 2015; Wu *et al*, 2001; Cordes *et al*, 1996). Using CRISPR genome engineering, we tagged Nup358 and Nup88 at their endogenous loci and examined their distribution in all tissues across *C. elegans* hermaphrodite development (Figure 1A). As expected, both Nups localized to the nuclear envelope in all cell types, including muscle, hypodermis, intestine, neurons, and germ cells (Figures 1B and S1A). In *C. elegans*, germ cell nuclei proliferate in a syncytial cytoplasm before individualizing to form sperm during the fourth larval (L4) stage and oocytes in adults (Figures 1A and S1B). We detected Nup358 and Nup88 in cytoplasmic foci in the residual body of spermatocytes, a transient structure that accumulates components discarded during spermatogenesis (Figure S1B). We also detected Nup358 and Nup88 in foci in growing oocytes and in early embryos (<∼80-cell stage) (Figures 1B, 1C, S1A, and S1C). We did not detect cytoplasmic Nup foci in somatic cells of young animals (Day 2 adults and younger). The cytoplasmic concentration of Nups in germ cells and early embryos was ∼3-5-fold higher than that observed in somatic cells (Figure 1D). We conclude that, in developing animals and young adults, Nup foci only form in gametes and early embryos, which accumulate higher levels of cytoplasmic Nups as compared to somatic tissues.

### Nup foci assembly is enhanced by oocyte arrest, heat stress, and aging

We noticed that the intensity of Nup foci in growing oocytes increased significantly between days 1 and 2 of adulthood (Figure S2A). Oocyte production occurs continuously in young hermaphrodites and slows down with age as sperm are depleted. To examine oocytes arrested in the absence of sperm, we used *fog-2(q71*) females which lack sperm and accumulate fully grown arrested oocytes in the oviduct (Schedl & Kimble, 1988). Strikingly, we observed a 14-fold increase in the intensity of Nup foci in the arrested oocytes of Day 1 adult *fog-2(q71*) females as compared to growing oocytes of age-matched wild-type hermaphrodites (Figures 2A and S2B). Previous studies have reported parallels between oocyte arrest and environmental stresses in inducing the formation of condensates in *C. elegans* oocytes (Jud *et al*, 2008; Elaswad *et al*, 2022). In agreement with these findings, we found that a 20-minute incubation at 30°C was sufficient to increase the intensity of Nup358 foci in growing oocytes by 16-fold (Figures 2A and S2B) and significantly increase Nup358 condensation in embryos (Figure 2B). In contrast, the same conditions of oocyte arrest and heat stress did not induce condensation of the stress granule scaffold G3BP (Figures 2A and S2B). Together these observations indicate that Nup foci assembly is readily enhanced by mild stress conditions. Consistent with this view, we found that >90% of Day 7 adult hermaphrodites exhibited Nup foci in somatic cells, including neurons and muscle cells (Figures 2C and D). We conclude that Nup foci are stress-sensitive structures that accumulate with age.

**Figure 2.**
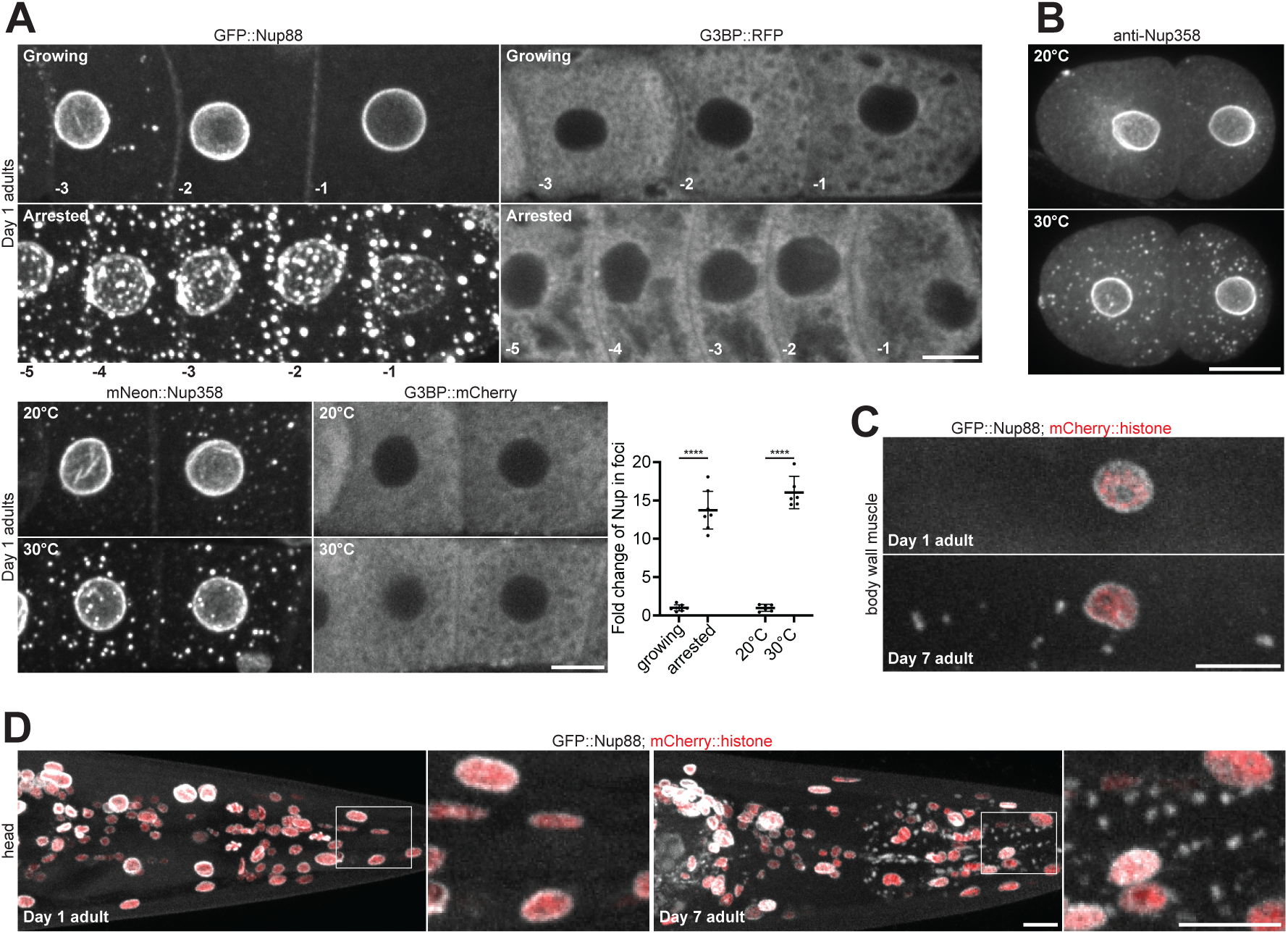
Cytoplasmic Nup foci increase with environmental stress and age. A. Top: Representative confocal micrographs showing CRISPR-tagged GFP::Nup88 in the proximal germline of Day 1 adults with growing (wild-type) versus arrested (*fog-2(q71)*) oocytes, or CRISPR-tagged G3BP::RFP in growing (mated) versus arrested (unmated) *fog-2(q71)* oocytes. Bottom left: Representative images showing CRISPR-tagged mNeonGreen::Nup358 and G3BP::mCherry in the -3 and -4 oocytes of Day 1 adults grown at 20°C or after shifting to 30°C for 20 min. Bottom right: Quantification of the percent of GFP::Nup88 in foci in growing versus arrested oocytes (n > 7 germlines) or mNeonGreen::Nup358 in foci at 20°C versus 30°C (n = 6 germlines). Values are normalized so that the average control condition (growing oocytes or 20°C) measurement = 1.0. See Figure S2B for raw (non-normalized) values of the percent Nup in foci for each condition. B. Representative confocal micrographs of endogenous Nup358 in 2-cell interphase embryos grown at 20°C or after shifting to 30°C for 20 min. C. Representative confocal micrographs showing GFP::Nup88 in body wall muscle cells of Day 1 versus Day 7 adults. Nuclei are marked by a mCherry::histone transgene. D. Representative confocal micrographs showing GFP::Nup88 in the head of a Day 1 versus Day 7 adult. Nuclei are marked by a mCherry::histone transgene. Areas indicated by white boxes are magnified at right. 100% (n = 11) of Day 1 adults lacked foci in somatic cells whereas cytoplasmic foci were observed outside of the germline in 92% (n = 12) of Day 7 adults. ****, P<0.0001. All images in this figure are maximum intensity projections, with the exception of G3BP (panel A) which are single focal planes. Scale bars = 10 μm.

### Nup foci in oocytes only contain FG-Nups and their direct binding partners

Nup foci in oocytes have been proposed to correspond to 1) condensates containing pore assembly intermediates or to 2) fully formed pore complexes in membranous annulate lamellae (Hampoelz *et al*, 2019b). To systematically compare the composition and stoichiometry of Nup foci to that of mature nuclear pore complexes at the nuclear envelope, we used a collection of genomically-encoded tags, transgenes, and antibodies against 16 Nups (including representatives of each nuclear pore subcomplex) as well as the Nup358 binding partners RanGAP and NXF1 (Figure 3A and Table S1). We examined Nup distribution in growing oocytes of Day 2 adult wild-type hermaphrodites where Nup foci are prominent.

**Figure 3.**
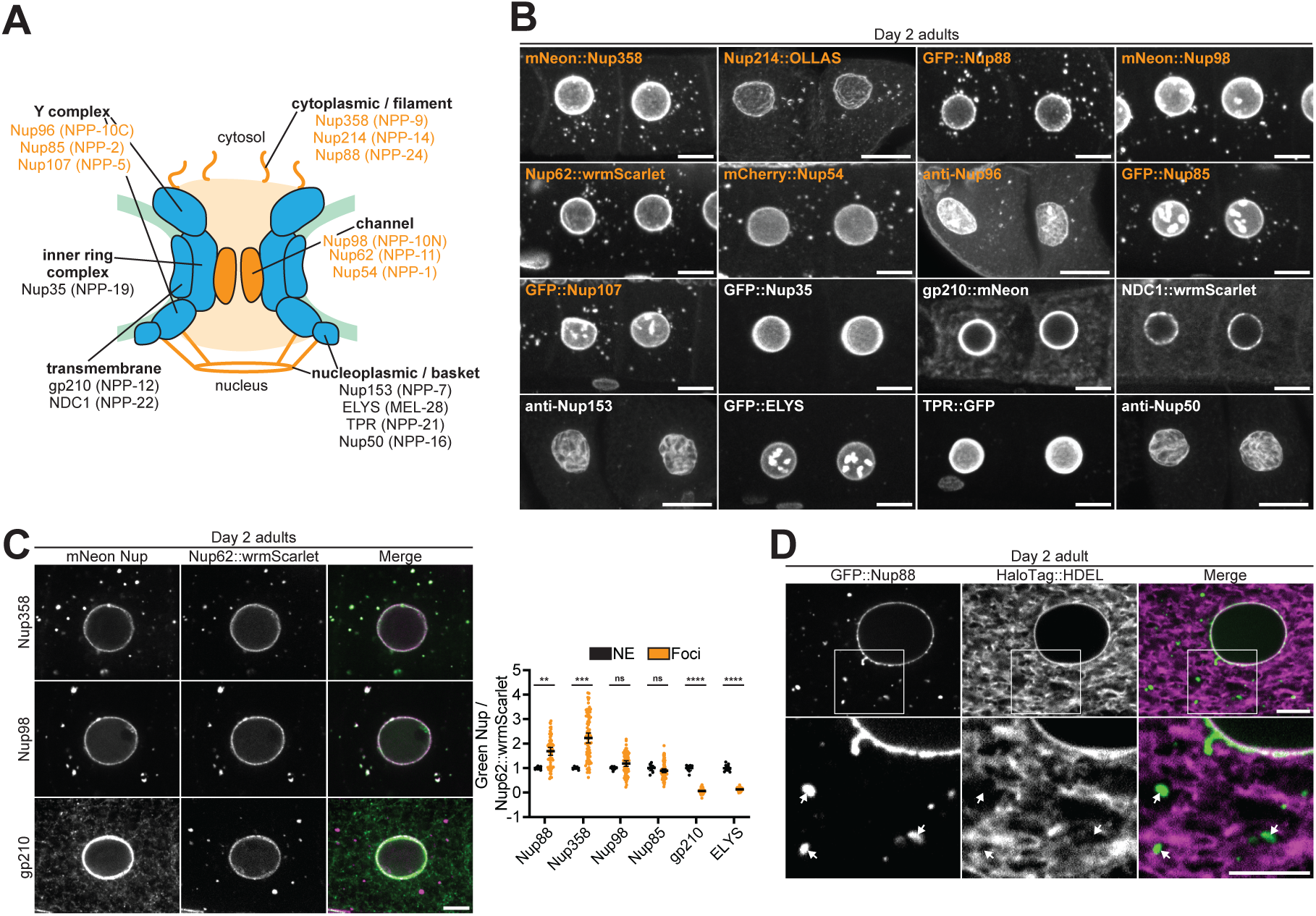
Cytoplasmic Nup foci contain only FG-Nups and their binding partners. A. Schematic depicting the location of all visualized Nups within the nuclear pore complex. Nups listed in orange localize to cytoplasmic foci in oocytes, whereas those denoted in black do not. Nups are listed with human names, with *C. elegans* homologs in parentheses. B. Representative confocal micrographs of the -3 and -4 oocytes from Day 2 adult *C. elegans* expressing tagged versions of each indicated Nup, or stained with antibodies against endogenous Nups. All images are maximum intensity projections, with the exception of gp210 and NDC1 which are single imaging planes. Orange labels designate Nups enriched in cytoplasmic foci. C. Left: Representative confocal micrographs of Day 2 adult oocytes showing colocalization of CRISPR-tagged Nup62::wrmScarlet with mNeonGreen-tagged Nup358, Nup98, or gp210. Nup358 represents a Nup that is enriched in foci relative to the nuclear envelope, whereas gp210 is absent from foci. Right: Quantification of the overlap between Nup62::wrmScarlet and each indicated Nup at the nuclear envelope (NE) versus cytoplasmic foci. Each point designates an individual nucleus or focus. Values are normalized so that the average ratio at the nuclear envelope = 1.0. Error bars represent 95% CI for n > 7 (nuclei) or n > 59 (foci). D. Representative confocal micrographs showing overlap of CRISPR-tagged GFP::Nup88 with the luminal endoplasmic reticulum/nuclear envelope marker HaloTag::HDEL in a Day 2 adult oocyte. 20% of foci completely overlapped with HaloTag::HDEL, 64% partially overlapped, and 16% showed no overlap with HaloTag::HDEL (n = 118). Areas indicated by white boxes are magnified below; white arrows indicate foci that do not completely overlap with the endoplasmic reticulum. ****, P<0.0001; ***, P<0.001; **, P<0.01; ns, not significant. Scale bars = 10 μm (panel B) or 5 μm (panels C and D).

As expected, all Nups tested localized to the nuclear envelope (Figures 3B and S3A). Nuclear basket and Y complex Nups additionally localized to the nucleoplasm and meiotic chromosomes, respectively, as previously described (Hattersley *et al*, 2016). Surprisingly, only a subset of Nups localized to cytoplasmic foci, including FG-Nups of the central channel and cytoplasmic filaments (Nup62, Nup98, Nup214, and Nup358) and their binding partners (Y complex Nups, Nup88, RanGAP, and NXF1) (Figures 3B, S3A, and S3B). The transmembrane Nups gp210 and NCD1 could be detected throughout the endoplasmic reticulum as previously described (Galy *et al*, 2008), but did not enrich in foci, nor did Nup35, an inner ring complex Nup. All nucleoplasmic-facing Nups (Nup153, Nup50, TPR, and ELYS) were enriched in the nucleoplasm and absent from cytoplasmic foci. We also analyzed the distribution of Nups in 4-cell stage early embryos and obtained the same results except for Nup35, which did not localize to foci in oocytes but did in embryos (Figures S3C and D). We conclude that cytoplasmic Nup foci contain only a subset of Nups and are primarily enriched for cytoplasm-facing FG-Nups and their binding partners (Figure 3A).

Co-staining experiments using the mAb414 antibody (Davis & Blobel, 1986) suggested that Nup foci contain multiple Nups (Figures S3A and D). To examine Nup stoichiometry in the foci, we crossed a subset of GFP-tagged Nups pairwise with Nup62::wrmScarlet. As expected, all Nups tested colocalized with Nup62::wrmScarlet at the nuclear envelope (Figure 3C). Nups that localize to cytoplasmic foci (Nup85, Nup88, Nup98, and Nup358) additionally colocalized with Nup62::wrmScarlet in all foci. Quantification of the ratio of the GFP-tagged Nup to Nup62::wrmScarlet revealed that each Nup accumulates in fixed stoichiometry relative to Nup62 at the nuclear envelope. In contrast, Nups exhibited variable stoichiometry in the cytoplasmic foci (Figure 3C).

Annulate lamellae pore complexes assemble on endoplasmic reticulum membranes (Stafstrom & Staehelin, 1984; Cordes *et al*, 1996) and have been observed in ∼10% of arrested *C. elegans* oocytes but not in growing oocytes or embryos (Langerak *et al*, 2019; Patterson *et al*, 2011; Pitt *et al*, 2000). We found that 80% of GFP::Nup88 foci in growing oocytes and 58% in arrested oocytes did not fully overlap with a marker for endoplasmic reticulum membranes (Figures 3D and S3E). We conclude that the majority of Nup foci are unlikely to correspond to nuclear pore assembly intermediates or annulate lamellae, as they lack critical nuclear pore scaffolds, exhibit variable Nup stoichiometry, and do not always associate with endoplasmic reticulum membranes.

### Nup foci are condensates scaffolded by excess FG-Nups

*In vitro*, FG-Nups readily condense into hydrogels (Labokha *et al*, 2012) raising the possibility that cytoplasmic Nup foci might form by spontaneous condensation of FG-Nups in the saturated environment of the oocyte. Condensation is highly sensitive to concentration: proteins de-mix into dense and dilute phases when their concentration exceeds the saturation concentration (*C_sat_*), the maximum concentration allowed in the soluble, dilute phase (Alberti *et al*, 2019).

To estimate the percent of Nup molecules that undergo condensation, we used Imaris software to quantify Nup fluorescence in nuclei, the cytoplasm, and cytoplasmic foci (Figure S4A and see materials and methods). Remarkably, we found that the vast majority of Nups distribute between a nuclear pool (∼30-40%) and a diffuse cytoplasmic pool (∼60-70%), with less than 3% of Nup molecules in foci (Figure 4A). The soluble cytoplasmic pool is the least concentrated but largest by volume and is readily visualized in sum projection micrographs (Figure S4B). These observations suggest that FG-Nups are maintained in oocytes at concentrations just in excess of the saturation concentration, such that most molecules are soluble and only a minority condense in the foci. If so, we predicted that removal of individual FG-Nups may be sufficient to drop below the threshold for condensation and reduce foci formation.

**Figure 4.**
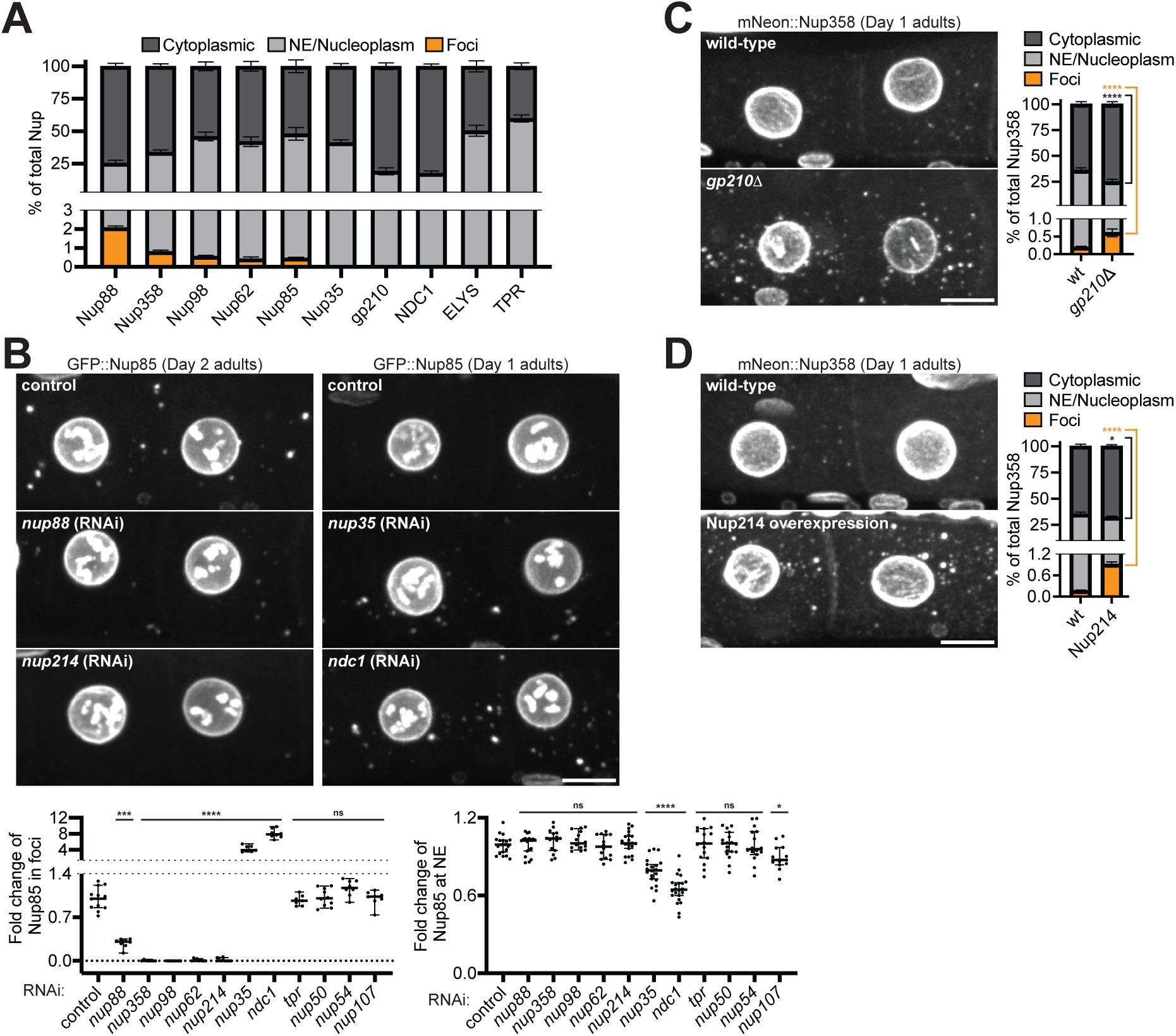
Nup foci are condensates scaffolded by cytoplasmic facing FG-Nups. A. Quantification of the distribution of each designated CRISPR-tagged Nup between the cytoplasm (soluble), nuclear envelope (NE)/nucleoplasm, and cytoplasmic foci. Measurements were made using the -3 and -4 oocytes of Day 2 adults. Error bars represent 95% CI for n > 5 germlines. B. Top: Representative confocal micrographs showing -3 and -4 oocytes of Day 1 or Day 2 adults with CRISPR-tagged GFP::Nup85. *nup214* RNAi is representative of a treatment that largely abolishes Nup foci, whereas Nup foci were partially diminished following RNAi-mediated knockdown of Nup88. Both *nup35* and *ndc1* RNAi enhanced Nup foci. Bottom left: Quantification of the total percent of GFP::Nup85 in foci following each RNAi treatment. Values are normalized so that the average control measurement = 1.0. Error bars represent 95% CI for n > 7 germlines. Bottom right: Line-scan quantification measuring GFP::Nup85 signal at the NE following each RNAi treatment. Values are normalized so that the average control measurement = 1.0. Error bars represent 95% CI for n > 13 nuclei. C. Left: Representative confocal micrographs showing CRISPR-tagged mNeonGreen::Nup358 in -3 and -4 oocytes of Day 1 wild-type versus *gp210Δ* adults. Right: Quantification of the distribution of mNeonGreen::Nup358 between the cytoplasm, NE/nucleoplasm, and cytoplasmic foci in wild-type versus *gp210Δ* oocytes. Error bars represent 95% CI for n > 6 germlines. D. Left: Representative confocal micrographs showing mNeonGreen::Nup358 in -3 and -4 oocytes of Day 1 adults with or without overexpression of Nup214::wrmScarlet. Right: Quantification of the distribution of mNeonGreen::Nup358 between the cytoplasm, NE/nucleoplasm, and cytoplasmic foci in wild-type oocytes versus those with Nup214 overexpression. Error bars represent 95% CI for n > 9 germlines. ****, P<0.0001; ***, P<0.001; *, P<0.05; ns, not significant. All images in this figure are maximum intensity projections. Scale bars = 10 μm.

We used RNAi and mutagenesis to systematically deplete individual Nups and examined the effect on Nup foci formation. As expected, depletion of non-FG or nucleoplasmic Nups, which are not present in foci, had no effect on foci formation (Figures 4B and S4C-E). In contrast, depletion of individual cytoplasm-facing FG-Nups (Nup62, Nup98, Nup214, or Nup358) essentially abolished the formation of Nup foci without affecting Nup levels at the nuclear envelope. Depletion of Nup88, which is structured but interacts with multiple subcomplexes containing FG-Nups (Fornerod *et al*, 1997; Griffis *et al*, 2003; Xylourgidis *et al*, 2006; Yoshida *et al*, 2011), partially depleted Nup foci, suggesting that interactions among FG-Nup subcomplexes contribute to foci formation. Loss of Nup35 or the transmembrane Nups NDC1 or gp210, in contrast, enhanced foci formation in oocytes, and also induced the formation of ectopic foci in the syncytial cytoplasm of the distal germline (Figures 4B, 4C, S4C, and S4F). As expected for structural Nups (Mansfeld *et al*, 2006; Mauro *et al*, 2022; Ródenas *et al*, 2009), depletion of NDC1 or Nup35 also decreased Nup levels at the nuclear envelope. We conclude that Nup foci assembly in oocytes depends primarily on the cumulative effect of high concentrations of the FG-Nups Nup62, Nup98, Nup214, and Nup358 in the cytoplasm.

To test whether high levels of FG-Nups are sufficient to drive foci formation, we generated a transgenic strain with an extra copy of *nup214::wrmScarlet* expressed under the control of the germline-specific *mex-5* promoter (Fan *et al*, 2020). We found that overexpression of Nup214::wrmScarlet was sufficient to increase the proportion of endogenous mNeonGreen::Nup358 in Nup foci by 4-fold (Figures 4D and S4G).

FG-Nup hydrogels assembled *in vitro* are readily dissolved by the aliphatic alcohol 1,6-hexanediol (Schmidt & Görlich, 2015), which disrupts hydrophobic interactions and has been reported to dissolve Nup foci in yeast, *Drosophila*, and HeLa cells (Hampoelz *et al*, 2019b; Patel *et al*, 2007; Agote-Aran *et al*, 2020). As expected, we found that hexanediol treatment dissolved Nup foci in *C. elegans*, although it had no effect on Nups at the nuclear envelope (Figure S4H). We conclude that Nup foci are FG-Nup condensates that arise when the cytoplasmic concentration of FG-Nups exceeds the saturation concentration.

### Nup foci disassemble during the oocyte-to-embryo transition and are not required for nuclear pore assembly in embryos

Although Nup foci are readily apparent in growing and arrested oocytes, we found that they were absent from oocytes that have entered meiotic M phase immediately prior to fertilization (−1 position in the oviduct; Figure 5A). Both the cytoplasmic concentration and total levels of Nup were higher in maturing oocytes as compared to growing oocytes (Figures 5A and S5A), indicating that Nup solubility increases during maturation. Similarly, Nup foci in early embryos disassembled at each mitosis and reassembled in interphase (Figure 5B and Videos S1 and 2), while overall Nup levels remained unchanged (Figure S5B). We conclude that the solubility of FG-Nups increases during M phase, causing the dissolution of Nup foci. These observations also indicate that Nup foci are transient structures that are not maintained during the oocyte-to-embryo transition and therefore are unlikely to contribute pre-assembled pore complexes for nuclear growth during embryogenesis.

**Figure 5.**
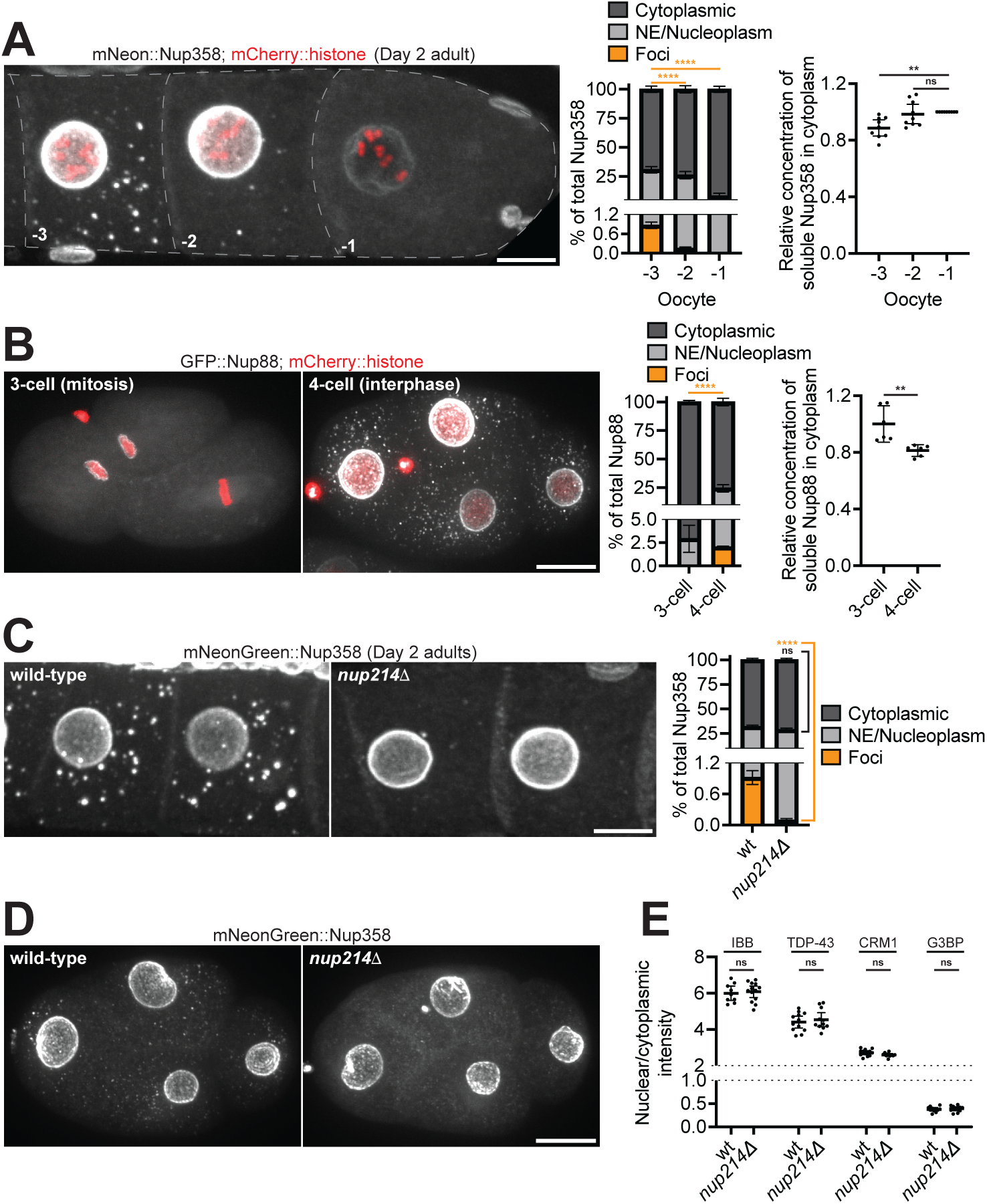
Nup foci are transient, non-essential condensates. A. Left: Representative confocal micrograph showing CRISPR-tagged mNeonGreen::Nup358 in Day 2 adult oocytes. Nup foci accumulate progressively throughout oocyte growth and peak in the -3 and -4 oocytes. Foci begin to disassemble in the -2 oocyte, and are absent from the -1 oocyte coincident with nuclear envelope breakdown. Middle: Quantification of the distribution of mNeonGreen::Nup358 between the cytoplasm (soluble), nuclear envelope (NE)/nucleoplasm, and cytoplasmic foci. Error bars represent 95% CI for n = 8 germlines. Right: Cytoplasmic (soluble) mNeonGreen::Nup358 fluorescence normalized to volume in -1, -2, and -3 oocytes. Values are normalized within the same germline so that the -1 oocyte measurement = 1.0. Error bars represent 95% CI for n = 9 germlines. B. Left: Representative confocal micrographs of CRISPR-tagged GFP::Nup88 in 3-cell (mitosis) versus 4-cell (interphase) embryos. Middle: Quantification of the distribution of GFP::Nup88 between the cytoplasm, NE/nucleoplasm, and cytoplasmic foci. Error bars represent 95% CI for n > 6 embryos. Right: Cytoplasmic (soluble) GFP::Nup88 fluorescence normalized to volume in 3-cell mitotic embryos versus 4-cell interphase embryos. Values are normalized so that the average 3-cell embryo measurement = 1.0. Error bars represent 95% CI for n > 6 embryos. C. Left: Representative confocal micrographs showing mNeonGreen::Nup358 in -3 and -4 oocytes of wild-type versus *nup214Δ* Day 2 adults. Right: Quantification of the distribution of mNeonGreen::Nup358 between the cytoplasm (soluble), NE/nucleoplasm, and cytoplasmic foci in wild-type versus *nup214Δ* oocytes. Error bars represent 95% CI for n > 8 germlines. D. Representative confocal micrographs showing mNeonGreen::Nup358 in wild-type versus *nup214Δ* interphase 4-cell embryos. E. Quantification of the nuclear/cytoplasmic ratio of an IBB_domain_::mNeonGreen reporter or CRISPR-tagged TDP-43::wrmScarlet, CRM1::mNeonGreen, and G3BP::mCherry in 28-cell stage embryos. Values are normalized so that the average wild-type measurement = 1.0. Error bars represent 95% CI for n > 9 embryos (IBB_domain_::mNeonGreen), n > 11 embryos (TDP-43::wrmScarlet), n > 11 embryos (CRM1::mNeonGreen), or n = 11 embryos (G3BP::mCherry). ****, P<0.0001; **, P<0.01; ns, not significant. All images in this figure are maximum intensity projections. Scale bars = 10 μm.

To directly test whether Nup foci might contribute to pore assembly in embryos, we used CRISPR genome engineering to generate a complete deletion of the *nup214* locus. Consistent with our RNAi results, using three independent markers (mNeonGreen::Nup358, RanGAP::wrmScarlet, and mAb414), we found that Nup foci were greatly reduced in *nup214Δ* mutant oocytes and embryos (Figures 5C, 5D, S5C, and S5D). Despite lacking robust Nup foci, *nup214Δ* embryos were 100% viable (Figure S5E). Furthermore, nuclear pore formation was not disrupted in *nup214Δ* mutant embryos (Figure S5F), as evidenced by normal partitioning of cargos between the nucleus and cytoplasm, including the nuclear RNA-binding protein TDP-43 and an importin β binding (IBB) domain reporter (Lott & Cingolani, 2011) (Figure 5E). We conclude that Nup foci are non-essential structures that do not contribute significantly to nuclear pore assembly during embryogenesis.

### Multiple mechanisms enhance Nup solubility in the cytoplasm

The vast majority of FG-Nups in oocytes are maintained in a soluble cytoplasmic pool (Figure 4A). To identify mechanisms that promote Nup solubility, we first tested PLK1 and CDK1, two kinases active in oocytes and known to drive nuclear pore disassembly during nuclear envelope breakdown in M phase (Chase *et al*, 2000; De Souza *et al*, 2004; Huelgas-Morales & Greenstein, 2018; Laurell *et al*, 2011; Linder *et al*, 2017; Martino *et al*, 2017; Onischenko *et al*, 2005; Rahman *et al*, 2015; Kutay *et al*, 2021). Additionally, consistent with prior experiments in *Xenopus* oocytes (Beckhelling *et al*, 2003), we observed that CDK1 enriches in Nup foci in *C. elegans* oocytes (Figure S6A). We found that RNAi depletion of PLK1 and CDK1 increased the proportion of Nups in foci as well as at the nuclear envelope in growing oocytes (Figures 6A, 6B, S6B, and S6C). Inhibition of the phosphatase PP2A blocks nuclear pore complex and Nup foci assembly in *Drosophila* embryos (Onischenko *et al*, 2005). Similarly, we found that RNAi depletion of the scaffolding subunit of PP2A led to a striking loss of Nup foci as well as depletion of Nups from the nuclear envelope (Figures 6A, 6B, S6B, and S6C). We conclude that, in addition to regulating nuclear pore assembly, Nup phosphorylation by cell cycle kinases increases the solubility of Nups in the cytoplasm and that PP2A phosphatase activity counteracts this effect.

**Figure 6.**
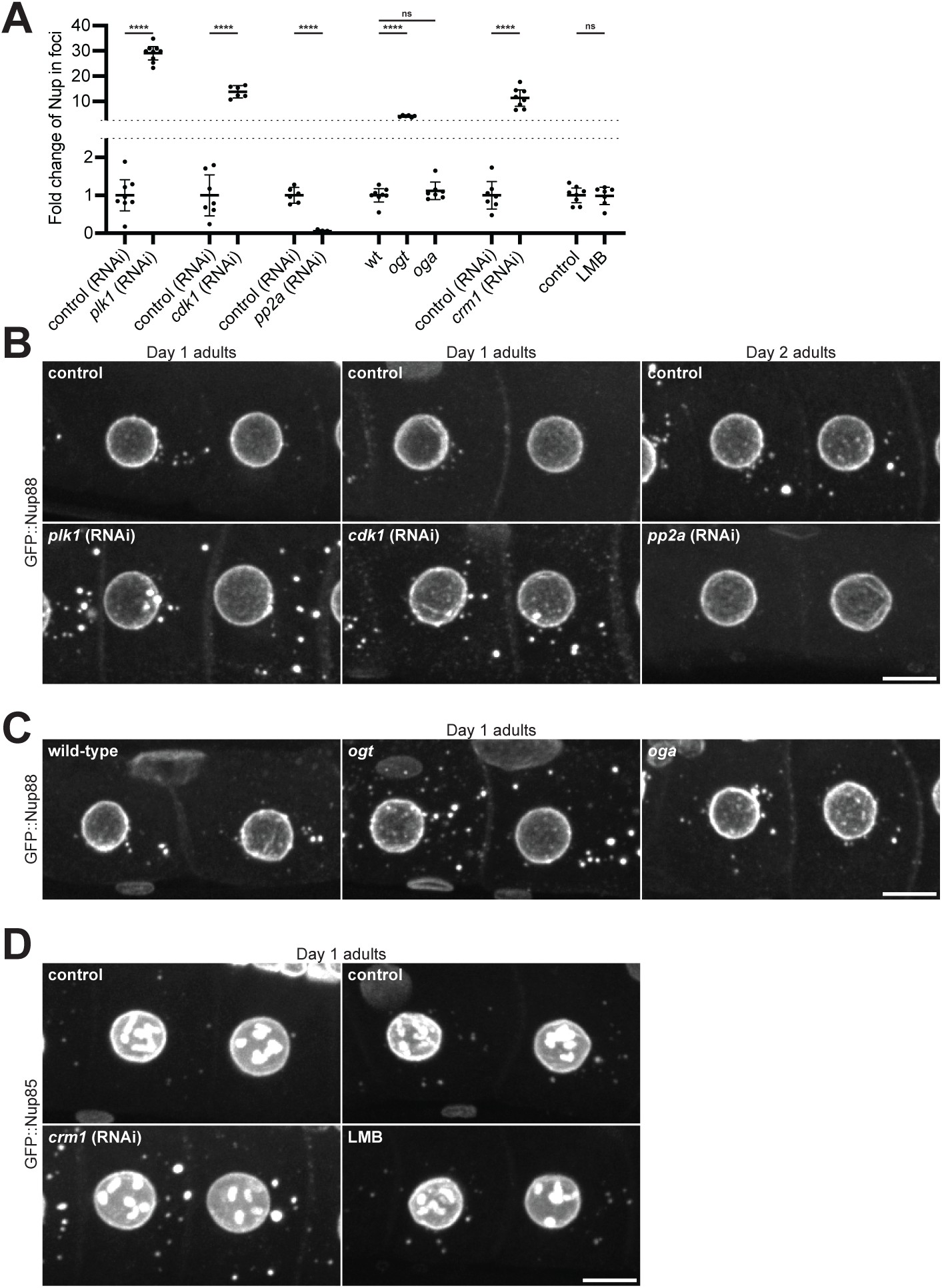
Phosphorylation, GlcNAcylation, and CRM1 promote Nup solubility. A. Compiled quantification of the percent of Nup in foci in each indicated condition. Values are normalized so that the average control condition measurement = 1.0. Data correspond to micrographs in Figure 6B (*plk1* RNAi, n > 8 germlines; *cdk1* RNAi, n > 6 germlines; *pp2A* RNAi, n > 6 germlines), Figure 6C (*ogt* and *oga* mutants, n > 6 germlines), and Figure 6D (*crm1* RNAi, n > 7 germlines; LMB treatment, n > 8 germlines). See Figure S6B for raw (non-normalized) values of the percent Nup in foci for each condition. B. Representative confocal micrographs showing CRISPR-tagged GFP::Nup88 in -3 and -4 oocytes depleted of PLK1, CDK1, or the PP2A scaffolding subunit PAA-1. Day 1 adults were used for kinase depletion, and Day 2 adults were used for phosphatase depletion. C. Representative confocal micrographs showing GFP::Nup88 in -3 and -4 oocytes of wild-type, *ogt*, or *oga* mutant Day 1 adults. D. Left: Representative confocal micrographs showing CRISPR-tagged GFP::Nup85 in -3 and -4 control oocytes or oocytes depleted of CRM1. Right: Representative confocal micrographs showing GFP::Nup85 in -3 and -4 oocytes of control animals or following treatment with the CRM1 inhibitor leptomycin B (LMB). All images are from Day 1 adults. ****, P<0.0001; ns, not significant. All images in this figure are maximum intensity projections. Scale bars = 10 μm.

Nup FG domains are heavily modified by O-GlcNAcylation (Ruba & Yang, 2016). The function of this modification remains unclear, but previous studies have proposed that O-GlcNAcylation may limit FG domain interactions within the central channel (Ruba & Yang, 2016; Yoo & Mitchison, 2021). O-GlcNAcylation is catalyzed by the enzyme O-GlcNAc transferase (OGT), which localizes to Nup foci in oocytes (Figure S6D). To test whether O-GlcNAcylation contributes to the solubility of cytoplasmic Nups, we measured foci formation in *ogt* mutant animals that lack Nup O-GlcNAcylation as previously described (Figures S6E and F) (Hanover *et al*, 2005). In support of a solubilizing role for O-GlcNAcylation, *ogt* mutants exhibited enhanced Nup foci formation (Figures 6A, 6C, S6B, and S6G). We also visualized Nup foci in a loss of function allele of the *C. elegans* O-GlcNAcase (OGA) reported to exhibit higher levels of Nup GlcNAcylation in embryos (Forsythe *et al*, 2006). We did not detect a significant change in Nup foci in the *oga* mutant, suggesting that, in oocytes, Nups may be sufficiently O-GlcNAcylated such that loss of OGA activity does not affect Nup solubility.

Recent studies have suggested that nuclear transport receptors (NTRs) function as chaperones to prevent aggregation of intrinsically disordered proteins (Guo *et al*, 2018; Hutten *et al*, 2020; Hofweber *et al*, 2018). We found that two endogenously tagged NTRs (CRM1 and transportin) localize to cytoplasmic Nup foci in *C. elegans* oocytes (Figures S7A and B). The exportin CRM1 makes high affinity interactions with the FG-Nup scaffolds Nup214 and Nup358 (Tan *et al*, 2018; Ritterhoff *et al*, 2016; Port *et al*, 2015). As we found Nup214 and Nup358 to be key scaffolds for Nup foci (see Figure 4B), we next tested whether CRM1 binding promotes the solubility of cytoplasmic Nups. Consistent with a solubilizing effect for CRM1 interaction, RNAi depletion of CRM1 led to an increase in Nup foci formation (Figures 6A, 6D, S6B, S7C, and S7D).

This effect is unlikely to be due to impaired nuclear export, as Nup foci were not altered in worms treated for 4 hours with the CRM1 inhibitor leptomycin B (LMB) (Figures 6A, 6D, S6B, S7D, and S7E). RNAi depletion of transportin did not affect Nup solubility (Figures S7F and G), indicating that CRM1 may be uniquely effective at solubilizing cytoplasmic Nups in oocytes. In summary, we conclude that Nup solubility in the cytoplasm is enhanced by multiple mechanisms including phosphorylation, O-GlcNAcylation, and CRM1 binding, which limit the formation of Nup foci.

### Ectopic Nup condensation in neurons is toxic

We only detected Nup foci in somatic cells in aged hermaphrodites (see Figures 2C and D). To determine whether Nup condensation might be detrimental in somatic cells, we used the neuron-specific *rab-3* promoter to overexpress Nup98, a highly cohesive FG-Nup that interacts with multiple structured Nups (Onischenko *et al*, 2017; Schmidt & Görlich, 2016). Unlike Nup98 expressed from its endogenous locus, overexpressed Nup98 readily formed cytoplasmic foci (Figure S8A). Remarkably, the ectopic Nup98 foci recruited endogenous Nup62, resulting in partial depletion of Nup62 from the nuclear envelope (Figure 7A). In control, non-neuronal cells that did not express the Nup98 transgene, Nup62 localized to the nuclear envelope as in wild-type animals (Figure 7B). Consistent with disrupted nuclear transport in the Nup98 overexpressing neurons, the nuclear protein TDP-43 was mislocalized to the cytoplasm (Figure S8B). Strikingly, *rab-3p*::Nup98 animals had shorter lifespans (Figure S8C) and appeared uncoordinated (barely moving) on plates or in liquid (Figure 7C and Videos S3 and 4), consistent with neuronal dysfunction and paralysis (Dimitriadi & Hart, 2010). We conclude that uncontrolled Nup condensation in neurons is toxic and leads to cellular dysfunction.

**Figure 7.**
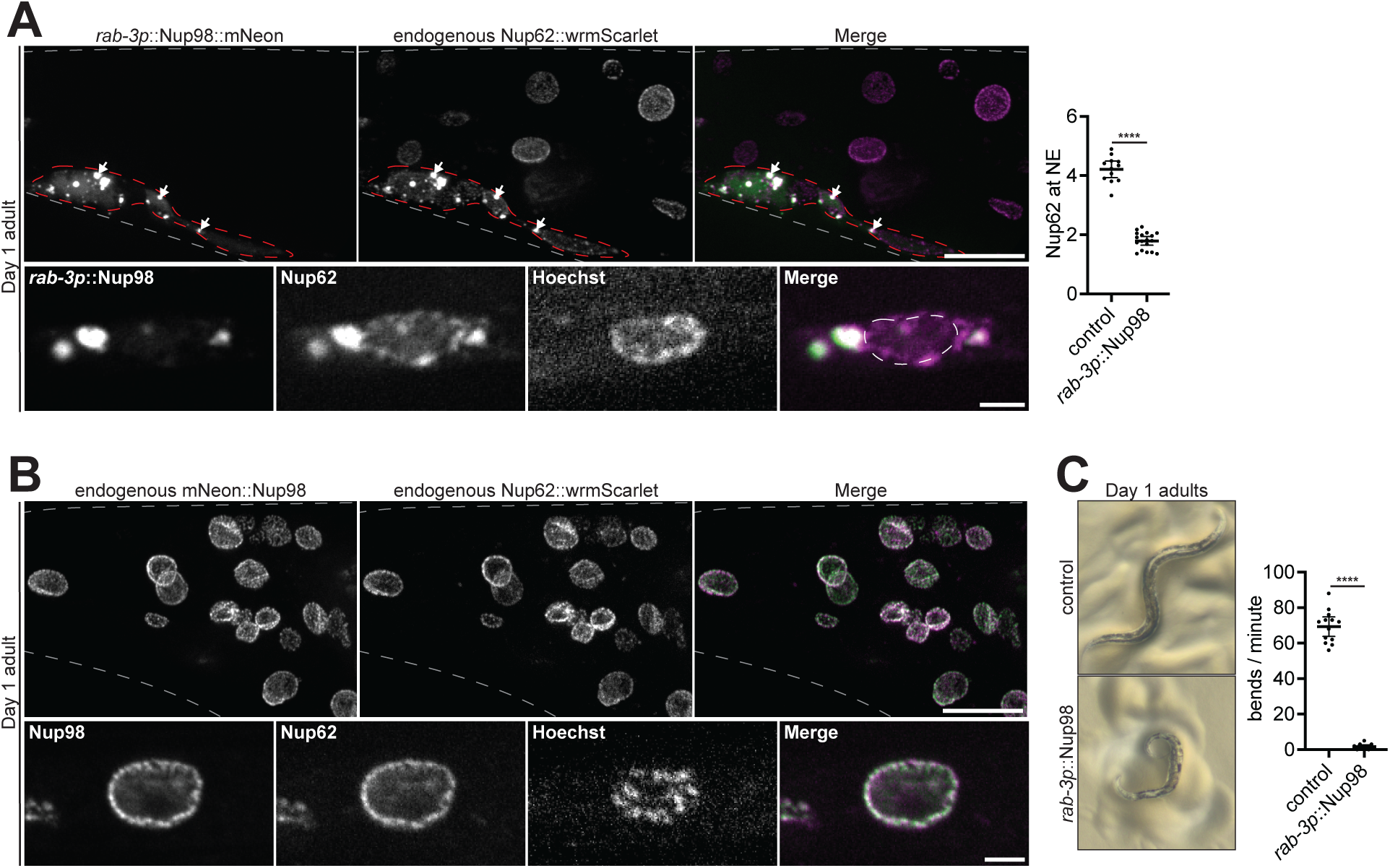
Ectopic Nup98 foci deplete endogenous Nups from the nuclear envelope and cause paralysis. A. Top left: Representative confocal micrographs showing colocalization of endogenous CRISPR-tagged Nup62::wrmScarlet with transgenic *rab-3p*::Nup98::mNeonGreen in the tail of a Day 1 adult. Gray dashed lines indicate the boundary of the tail. Note that transgenic *rab-3p*::Nup98::mNeonGreen is only expressed in neurons, which are designated by the red dashed outline. White arrows indicate colocalization of endogenous Nup62 with ectopic Nup98 foci. Bottom left: Representative confocal micrographs showing endogenous Nup62 depletion from the nuclear envelope. White dashed lines indicate the boundary of the nucleus. Right: Line-scan quantification of the nuclear envelope (NE) to nucleoplasm ratio of endogenous Nup62 in control (non-neuronal) cells, versus neurons with ectopically expressed *rab-* 3p::Nup98::mNeonGreen. Error bars represent 95% CI for n > 12 nuclei. B. Top: Representative confocal micrographs showing colocalization of Nup62::wrmScarlet with CRISPR-tagged endogenous mNeonGreen::Nup98 in the tail of a Day 1 adult. Gray dashed lines indicate the boundary of the tail. Bottom: Representative confocal micrographs showing colocalization of Nup62 with Nup98 at a single nucleus. C. Left: Representative images of a control Day 1 adult versus a Day 1 adult expressing Nup98::mNeonGreen driven by the pan-neuronal *rab-3* promoter. Right: Quantification of the swimming behavior of control Day 1 adults versus those with ectopically expressed *rab-3p*::Nup98::mNeonGreen. Error bars represent 95% CI for n > 11 worms. ****, P<0.0001. All images in this figure are maximum intensity projections, with the exception of panels A and B (bottom) which are single focal planes. Scale bars = 2 μm (panels A and B, bottom) or 10 μm (panels A and B, top).

## Discussion

Cytoplasmic Nup foci have been observed in oocytes, yeast, and in many cell types in culture (Cordes *et al*, 1996; Raghunayakula *et al*, 2015; Ren *et al*, 2019; Colombi *et al*, 2013). In this study, we report the first systematic examination of the incidence of Nup foci across all tissues in an intact animal. We find that Nup foci are rare in healthy animals and arise only in cells where cytoplasmic Nup concentration is highest: gametes and early embryos. Although Nup condensates appear prominent when observed by fluorescence microscopy, they account for less than 3% of total cellular Nups and consist primarily of highly cohesive FG-Nups. The vast majority of FG-Nups are stored as soluble molecules in the cytoplasm whose condensation is actively suppressed by multiple mechanisms. Stress and aging promote FG-Nup condensation which can be toxic if uncontrolled. Our findings do not support an essential role for Nup foci in pore assembly and suggest instead that Nup foci are accidental byproducts of the natural tendency of FG-Nups to condense.

### Cytoplasmic Nup foci arise by condensation of FG-Nups and their binding partners

In *C. elegans*, annulate lamellae form in a minority (∼10%) of arrested oocytes and are not observed in growing oocytes or embryos (Langerak *et al*, 2019; Pitt *et al*, 2000; Patterson *et al*, 2011). A recent study also reported rare co-localization of Nup160, a structural Nup, with the transmembrane Nup NDC1 in cytoplasmic puncta (Mauro *et al*, 2022). While these observations suggest that pre-pore-like structures occasionally form in the cytoplasm of *C. elegans* oocytes and embryos, as observed in other animals, our quantitative analyses clearly demonstrate that the majority of Nup foci are unlikely to correspond to annulate lamellae or other pore precursors, as they 1) lack Nups essential for pore assembly including transmembrane Nups, 2) account for less than 3% of total Nup molecules, 3) display heterogeneous Nup stoichiometry, and 4) do not colocalize with membranes. Consistent with a concentration-dependent assembly mechanism, depletion and overexpression of single FG-Nups eliminate and enhance, respectively, foci formation. Together these observations suggest that Nup foci are not structured pre-pore assemblies, but are condensates scaffolded by cohesive FG-Nups, including Nup62, Nup98, Nup214, and Nup358, and their binding partner Nup88.

Consistent with our findings, a recent systematic survey in HEK293T cells revealed that cytoplasm-facing FG-Nups and their binding partners accumulate in cytoplasmic foci, but Nup153, which faces the nucleoplasm, does not (Cho *et al*, 2022). Other studies in HeLa and Cos7 cells have also documented that most Nup foci do not colocalize with membranes (Ren *et al*, 2019; Agote-Aran *et al*, 2020). Similarly, Nup foci in yeast cells contain multiple FG-Nups but lack transmembrane or inner ring complex Nups (Colombi *et al*, 2013). In agreement with this study, we found that the FG-Nup Nup214 forms hexanediol-sensitive foci in yeast cells, but the nucleoplasm-facing Nups Nup50 and Nup153 do not (Figure S8D). Together, these studies suggest that, as we describe here for *C. elegans*, many reported Nup foci likely correspond to FG-Nup condensates rather than pre-assembled pore complexes.

We suggest that Nup foci arise whenever the concentration of FG-Nups exceeds the solubility threshold in the cytoplasm. Consistent with this hypothesis, depletion of scaffold nucleoporins that liberate FG-Nups enhance foci formation in *C. elegans* oocytes (Figures 4 and S4), yeast (Makio *et al*, 2009), and HeLa cells (Raghunayakula *et al*, 2015). Similarly, intranuclear Nup assemblies termed GLFG bodies were reported in HeLa cell lines with elevated levels of Nup98, a highly cohesive FG-Nup (Griffis *et al*, 2002; Morchoisne-Bolhy *et al*, 2015). Our findings indicate that, even under conditions where FG-Nups exceed their solubility limit by a small margin, they from bright foci easily visible by standard microscopy techniques.

### Phosphorylation, GlcNAcylation, and CRM1-mediated chaperoning suppresses Nup condensation

The majority of FG-Nup molecules in the cytoplasm are maintained in a soluble state. We have found that the same kinases that drive pore complex disassembly during M phase (PLK1 and CDK1) also promote Nup solubility in the cytoplasm during interphase. Other kinases implicated in Nup phosphorylation and pore disassembly, including NIMA and DYRK kinases (Laurell *et al*, 2011; De Souza *et al*, 2004; Wippich *et al*, 2013), could also contribute. Consistent with phosphorylation promoting Nup solubility, cellular fractionation experiments have shown that soluble Nups are more highly phosphorylated than Nups in pore complexes (Onischenko *et al*, 2004).

Consistent with prior findings showing that GlcNAcylated FG-Nups are less prone to condensation *in vitro* (Labokha *et al*, 2012; Schmidt & Görlich, 2015), we find that GlcNAcylation also contributes to Nup solubility in oocytes and embryos. Numerous studies have reported a protective role for O-GlcNAcylation in neurodegenerative disease (Lee *et al*, 2021), raising the possibility that this modification plays a general solubilizing role for aggregation-prone proteins.

Finally, we find that the nuclear export factor CRM1 also contributes to Nup solubility. Structural analyses of CRM1/Nup214 complexes reveal that hydrophobic patches on the surface of CRM1 make extensive contacts with Nup214 FG domains (Port *et al*, 2015). CRM1 generates high affinity interactions with both Nup214 and Nup358 that are significantly stronger than the weak, transient interactions characteristic of most Nup/NTR pairs (Tan *et al*, 2018; Ritterhoff *et al*, 2016; Port *et al*, 2015). Both Nup214 and Nup358 are required for Nup foci formation and therefore neutralization of these proteins by CRM1 is predicted to reduce Nup foci formation. Interestingly, loss of another transport factor, transportin, did not affect Nup solubility in *C. elegans* oocytes, suggesting that in this cell type Nup solubility depends primarily on CRM1.

### Most Nup foci are unlikely to serve an essential biological role and are potentially toxic

In *Drosophila* oocytes, Nup condensates have been proposed to function as intermediates in the biogenesis of annulate lamellae that supply pre-made nuclear pores for use during embryogenesis (Hampoelz *et al*, 2019b). Our analyses do not support such a role for Nup foci in *C. elegans*. First, Nup foci only account for a small proportion (<3%) of total cellular Nups, as also observed in *Drosophila* embryos (Onischenko *et al*, 2004) and egg chambers (Hampoelz *et al*, 2019b). Second, Nup foci are transient structures that dissolve fully at oocyte maturation and every M phase thereafter. Third, we generated a mutant that severely reduces the incidence of Nup foci in oocytes and embryos yet assembles functional nuclear pores during embryogenesis and is 100% viable. In *Drosophila* egg chambers, Nup condensates assemble in nurse cells and are transported into the oocyte by a microtubule-dependent mechanism required for the formation of annulate lamellae (Hampoelz *et al*, 2019b). These observations suggest that, in that system, Nup condensation has been harnessed as a mechanism to concentrate Nups in oocytes. The extent to which annulate lamellae assembled in oocytes contribute to nuclear pores in embryos, however, remains to be determined as the vast majority of Nups are soluble in *Drosophila* embryos (Onischenko *et al*, 2004), as we show here for *C. elegans*.

We suggest that, in most cell types, Nup foci are non-essential byproducts of FG-Nup condensation that are potentially stress-induced and toxic. Consistent with our findings, essential functions have not been identified for Nup foci observed in cell culture, including GLFG bodies and CyPNs (cytoplasmic accumulations of PML and nucleoporins)(Jul-Larsen *et al*, 2009; Griffis *et al*, 2002). In contrast, several studies suggest connections to stress and disease, including 1) Nup accumulation in stress granules (Zhang *et al*, 2018; Agote-Aran *et al*, 2020), 2) aberrant condensation of Nup98 and Nup214 fusion proteins driving oncogenic transformation in certain leukemias (Zhou & Yang, 2014; Chandra *et al*, 2022; Terlecki-Zaniewicz *et al*, 2021), 3) the formation of nuclear envelope associated Nup condensates in models of DYT1 dystonia (Prophet *et al*, 2022), and 4) the presence of Nups in pathological inclusions in primary patient samples and models of neurodegenerative disease (Chandra & Lusk, 2022; Fallini *et al*, 2020; Hutten & Dormann, 2020). We find that overexpression of Nup98 in neurons initiates the formation of toxic condensates that cause neuronal dysfunction. Our findings are also consistent with recent studies reporting that cytoplasmic FG-Nups drive aggregation of TDP-43 in both ALS/FTLD and following traumatic brain injury (Anderson *et al*, 2021; Gleixner *et al*, 2022).

The deleterious effects of Nup condensation are likely context dependent. In arrested oocytes, Nup condensation increases ∼14-fold over growing oocytes, yet is likely not damaging as the majority of arrested oocytes go on to form viable embryos when fertilized (Jud *et al*, 2008). Pore complexes and Nup condensates in oocytes and embryos fully disassemble during M phase, allowing for a cycle of “renewal” with each cell division. We suggest that Nup condensation is only dangerous in post-mitotic cells that lack M phase-specific Nup solubilizers and where certain Nups are naturally long-lived (D’Angelo *et al*, 2009; Toyama *et al*, 2013). Somatic cells likely avoid Nup condensation primarily by maintaining low levels of cytoplasmic Nups relative to gametes. We did, however, observe Nup foci in the somatic cells of aged adults, which may be linked to the progressive decline in proteostasis that initiates during *C. elegans* adulthood (Herndon *et al*, 2002; Ben-Zvi *et al*, 2009). It will be interesting to investigate whether the mechanisms that dissolve Nup condensates in oocytes could be used to reverse Nup condensation in aged somatic tissues.

## Materials and methods

### *C. elegans* and yeast strains and culture

*C. elegans* were cultured using standard methods (Brenner, 1974). Briefly, worms were maintained at 20°C on normal nematode growth media (NNGM) plates (IPM Scientific Inc. cat # 11006-548) seeded with OP50 bacteria. We have found that Nup solubility is highly influenced by multiple factors including animal age: for all Nups tested the number and size of oocyte foci increased significantly between days 1 and 2 of adulthood (see Figure S2A). Therefore, for all experiments worms were synchronized as Day 1 or 2 adults using vulval morphology to stage L4 larvae. The age of animals used for each experiment is indicated in figures and legends.

Endogenous *npp-21* (TPR) was tagged with GFP using CRISPR/Cas9-mediated genome editing as previously described (Arribere *et al*, 2014). Endogenous *npp-24* (Nup88) and *npp-2* (Nup85) were tagged with G>F>P using SapTrap CRISPR/Cas9 gene modification as previously described (Schwartz & Jorgensen, 2016). G>F>P contains Frt sites in introns 1 and 2 of GFP that enable FLP-mediated, conditional knockout; in the absence of FLP, the construct behaves as normal GFP. Endogenous *npp-19* (Nup35) was tagged with G>F>P based on protocols for nested CRISPR (Vicencio *et al*, 2019) and “hybrid” partially single-stranded DNA donors (Dokshin *et al*, 2018). All other endogenous edits were performed using CRISPR/Cas9-mediated genome editing as described previously (Paix *et al*, 2017). Transgenic Nup214 and Nup98 strains (JH4119, JH4204, and JH4205) were generated using SapTrap cloned vectors as previously described (Fan *et al*, 2020). Standard crosses were used to generate strains with multiple genomic edits. All strains used or generated in this study are described in Table S1.

Yeast strains were generated using homologous recombination of PCR-amplified cassettes (Longtine *et al*, 1998). Endogenous *NUP159* (Nup214), *NUP60* (Nup153), and *NUP2* (Nup50) were tagged by amplifying the *mNeonGreen::HIS3* cassette from pFA6a-mNeonGreen::HIS3 (Thomas *et al*, 2019) using primers with homology to the C-termini (without the stop codon) and downstream regions of the genes. Yeast strains generated in this study are described in Table S1.

### RNAi

RNAi was performed by feeding (Timmons & Fire, 1998). RNAi vectors were obtained from the Ahringer or Open Biosystems libraries and sequence verified, or alternatively cloned from *C. elegans* cDNA and inserted into the T777T enhanced RNAi vector (Addgene cat # 113082). RNAi feeding vectors were freshly transformed into HT115 bacteria, grown to log phase in LB + 100 μg/mL ampicillin at 37°C, induced with 5 mM IPTG for 45 min, and plated on RNAi plates (50 μg/mL Carb, 1 mM IPTG; IPM Scientific Inc. cat # 11006-529). Seeded plates were allowed to dry overnight at RT before adding L4 larvae or Day 1 adults. For depletion of Nup98 (Figures 4B and S4C), RNAi feeding was performed for 6 hr at 25°C; partial depletion was used to minimize cytological defects caused by loss of Nup98. For all other experiments, RNAi feeding was performed for 18-24 hr at 25°C. For all experiments, control worms were fed HT115 bacteria transformed with the corresponding L4440 or T777T empty vector.

### Immunofluorescence

For immunostaining of embryos, gravid adults were placed into 7 μL of M9 media on a poly-L-lysine coated slide and compressed with a coverslip to extrude embryos. For immunostaining of oocytes, staged adults were dissected on poly-L-lysine slides to extrude the germline, and a coverslip was placed gently on top. In both cases, slides were immediately frozen on aluminum blocks pre-chilled with dry ice. After > 5 min, coverslips were removed to permeabilize embryos (freeze-cracking), and slides were fixed > 24 hr in pre-chilled MeOH at -20°C. Slides were then incubated in pre-chilled acetone for 10 min at -20°C, and blocked in PBS-T (PBS, 0.1% Triton X-100, 0.1% BSA) for > 30 min at RT. Slides were then incubated overnight in primary antibody in a humid chamber at 4°C. Slides were washed 3 x 10 min in PBS-T at RT, incubated in secondary antibody for 2 hr in a humid chamber at RT, and washed 3 x 10 min in PBS-T at RT. Slides were then washed 1x in PBS before being mounted using Prolong Glass Antifade Mountant with NucBlue (Thermo Fisher cat # P36981). Primary antibodies were diluted as follows: mAb414 (1:1,000; Biolegend cat # 902907), anti-Nup358 (1:250; Novus Biologicals cat # 48610002), anti-Nup50 (1:250, Novus Biologicals cat # 48590002), anti-GlcNAc RL2 (1:100; Invitrogen cat # MA1-072), anti-Nup96 (1:250, (Ródenas *et al*, 2012)), anti-Nup153 (1:250, (Galy *et al*, 2003)), anti-OLLAS-L2 (1:50, Novus Biologicals cat # NBP1-06713). Secondary antibodies were diluted as follows: Cy3 Donkey anti-Mouse IgG (1:200; Jackson cat # 715-165-151), AlexaFluor 488 Goat anti-Rabbit IgG (1:200; Invitrogen cat # A-11034), AlexaFluor 568 Goat anti-Rabbit IgG (1:200; Invitrogen cat # A-11011), Alexa Fluor 488 Goat anti-Rat IgG (1:200; Invitrogen cat # A-11006), AlexaFluor 488 anti-GFP (1:500; Invitrogen cat # A-21311).

### LMB and HXD treatment, HaloTag and Hoechst labeling, and heat stress

For CRM1 inhibition, leptomycin B (LMB; Sigma cat # L2913) was diluted in OP50 bacteria to a final concentration of 500 ng/mL and seeded on NNGM plates. 10-20 Day 1 adults were transferred to LMB or control vehicle plates and incubated at 20°C for 4 hr prior to imaging. For treatment of embryos with 1,6-hexanediol (HXD, Acros Organics cat # 629-11-8), L4 larvae were fed with *ptr-2* RNAi for 18-24 hr at 20°C. Embryos depleted of PTR-2, which permeabilizes the eggshell to allow for HXD treatment, were dissected into L-15 media (Thermo Fisher cat # 21083027) containing 2% HXD and immediately imaged. For HXD treatment of yeast, log-phase yeast were pelleted, re-suspended in media containing 5% HXD, and allowed to grow for 10 min at 30°C prior to imaging.

For HaloTag labeling, Janelia Fluor 646 HaloTag Ligand (Promega cat # GA1121) was diluted in OP50 bacteria to a final concentration of 25 μg/mL and seeded on NNGM plates. 10-20 L4 larvae or Day 1 adults were added and incubated without light at 20°C for 16-20 hr prior to imaging. For Hoechst staining, Hoechst 33342 dye (Thermo Fisher cat # 62249) was diluted in OP50 bacteria to a final concentration of 200 μM and seeded on NNGM plates. 10-20 L4 larvae were added and incubated without light at 20°C for 16-20 hr prior to imaging.

To induce heat stress, animals were grown at 20°C then transferred to pre-warmed NNGM plates at 30°C for 20 min prior to imaging at room temperature or processing for immunofluorescence.

### Embryonic viability and lifespan analysis

To measure embryonic viability, six Day 1 adults were transferred to six NNGM plates (36 animals total) and allowed to lay embryos for 1 hr at 20°C. Adults were then removed and the number of embryos on each plate was counted. Embryos were then allowed to hatch, and the number of adults on each plate was counted after 3 days at 20°C. Viability counts were repeated in three independent experiments, and embryonic viability was measured as the number of surviving adults divided by the original number of embryos counted in each experiment.

To measure adult lifespan, 75 Day 1 adults were transferred to five NNGM plates (15 animals per plate) and incubated at 20°C. Worms were scored daily and considered to be dead if they failed to move when prodded. Worms were transferred every 2 days to avoid progeny, and any worms that crawled off the plates were censored from analysis.

### Swimming assay

To measure swimming behavior, 5-10 Day 1 adults were transferred to a 33 mm culture dish (MatTek cat # P35G-1.5-14-C) containing 400 μL M9 media and immediately filmed using an Axiocam 208 color camera (Zeiss) mounted on a Stemi 508 Stereo Microscope (Zeiss). Swimming assays were performed at RT (∼22°C). Movies were exported to ImageJ, and the number of body bends per minute was counted manually.

### Imaging

For live imaging of germlines and somatic tissues, five staged adults were transferred to the middle well of a 3-chambered slide (Thermo Fisher cat # 30-2066A) in 10 μL of L-15 media with 1 mM levamisole. 20 μm polystyrene beads (Bangs Laboratories Inc. cat # PS07003) were then added to support a coverslip (Marienfeld cat # 0107052). Germlines were imaged using an inverted Zeiss Axio Observer with CSU-W1 SoRa spinning disk scan head (Yokogawa), 1x/2.8x/4x relay lens (Yokogawa), and an iXon Life 888 EMCCD camera (Andor) controlled by Slidebook 6.0 software (Intelligent Imaging Innovations). To image germlines or somatic cells, a 20 μm Z stack (1 μm step size) was captured using a 63x objective (Zeiss) with the 1x relay lens. For high resolution images of oocytes, 3 μm Z stacks (0.1 μm step size) were acquired using the 63x objective with the 2.8x relay lens. As germline condensates are highly sensitive to imaging-induced stress (Elaswad et al., 2022), care was taken to avoid compression of germlines, and all animals were imaged only once and maintained on the slide for <5 min. To image entire worms (Figure S8A), an 80 μm Z stack (1 μm step size) was captured using a 10x objective (Zeiss) with the 1x relay lens.

For live imaging of embryos, five young adults were transferred to 10 μL of L-15 media on a coverslip and dissected to release embryos. 20 μm polystyrene beads were then added to prevent compression, and the coverslip was inverted onto a microscope slide (Thermo Fisher cat # 12-550-403). Embryos were imaged as 15 μm Z stacks (1 μm step size), captured using the 63x objective with the 2.8x relay lens. For imaging fixed germlines and embryos, prepared slides were imaged as 15 μm Z stacks (0.5 μm step size), captured using the 63x objective with the 2.8x relay lens.

For live imaging of yeast, cells were grown overnight in synthetic dropout media (Thermo Fisher cat # DF0919-15-3) at 30°C and imaged in log-phase (OD_600_ of ∼0.5) at room temperature. Yeast were imaged as 6 μm Z stacks (0.5 μm step size), captured using the 63x objective with the 2.8x relay lens.

Images were exported from Slidebook software and further analyzed using ImageJ or Imaris image analysis software. For presentation in figures, images were processed using ImageJ, adjusting only the minimum/maximum brightness levels for clarity with identical leveling between all images in a figure panel. Images presented in figures are maximum intensity projections (10 μm for germlines, 15 μm for embryos, 6 μm for yeast) or single focal planes as indicated in the legends.

### Image quantification

The overlap of GFP or mNeonGreen-tagged Nups with Nup62::wrmScarlet (Figure 3C) was measured using single focal planes exported to ImageJ. The Nup62::wrmScarlet micrograph was used to create a mask defining the nuclear envelope as well as cytoplasmic foci as individual regions of interest (ROIs). This mask was then applied to both the GFP/mNeonGreen Nup micrograph as well as the Nup62::wrmScarlet micrograph and the integrated density was measured within each ROI. To control for cytoplasmic background, the average cytoplasmic signal for the GFP/mNeonGreen Nup was multiplied by the area of each ROI, and the resulting value subtracted from integrated density for the GFP/mNeonGreen Nup. Background normalized GFP/mNeonGreen Nup values were divided by Nup62::wrmScarlet values to obtain the ratio of GFP/mNeonGreen Nup to Nup62::wrmScalet at each ROI.

To quantify the overlap of GFP::Nup88 with membranes (Figures 3D and S3E), Z stacks of oocytes expressing GFP::Nup88 and the HaloTag::HDEL reporter were manually scored into 3 categories: 1. Complete overlap (the entire Nup88 focus overlapped with HaloTag::HDEL); 2. Partial overlap (the Nup88 focus partially overlapped or was directly adjacent to HaloTag::HDEL); 3. No overlap (the Nup88 focus did not directly contract membranes marked by HaloTag::HDEL).

To quantify the distribution of Nups in oocytes as well as total expression, Z stacks were exported to Imaris image analysis software. The “Surface” tool was first used to isolate the -3 and -4 oocytes from each germline (Figure S4A). For each pair of -3 and -4 oocytes, the Surface tool was then used to isolate both nuclei and the “Spot” tool was used to isolate cytoplasmic foci. The percent of Nup present at the nuclear envelope/nucleoplasm was measured as the intensity sum for both nuclei divided by the total intensity sum of the oocytes. Similarly, the percent of Nup present in foci was measured as the intensity sum for all foci divided by the total intensity sum of the oocytes. Finally, the percent soluble Nup was defined as 100% minus the percentage of Nup in both nuclei and foci. Total Nup expression was measured as the intensity sum of the -3 and -4 oocytes normalized to volume. To control for autofluorescent background in all measurements, staged animals lacking fluorescent tags were imaged using identical imaging settings. The average intensity sum per volume was calculated for the -3 and -4 oocytes of germlines lacking fluorescent tags and subtracted from the intensity sum measured for oocytes with tagged Nups. To measure the intensity of GFP::Nup35 per nuclear volume (Figure S5F), the intensity value for the -3 and -4 oocyte nuclei or 3 embryonic nuclei were divided by nuclear volume and the resulting values were averaged for each germline.

To quantify the distribution of Nups in embryos, Z stacks were exported to Imaris software. The Surface tool was used to isolate the entire embryo as well as all nuclei, and the Spot tool was used to isolate cytoplasmic foci. The percent of Nup at the nuclear envelope/nucleoplasm or foci was measured as the intensity sum of all nuclei or foci divided by the total intensity sum of the embryo, respectively. The percent soluble Nup was defined as 100% minus the percentage of Nup in nuclei and foci. For all measurements, embryos lacking fluorescent tags were used to control for autofluorescent background as described for oocytes.

The Y complex component Nup85 localizes to meiotic chromosomes and a high percentage of Nup85 is present in the nucleoplasm. Therefore, line-scan analysis was used to measure the amount of GFP::Nup85 at the nuclear envelope (Figure 4B). Z stacks were exported to ImageJ and line traces were drawn to pass through the central plane of -3 and -4 oocyte nuclei as well as the image background. For each nucleus, the two peak values of the nuclear envelope rim were averaged and normalized to the image background. Line-scan analysis was also used to quantify depletion of endogenous Nup62::wrmScarlet from the nuclear envelope in neurons expressing *rab-3p*::Nup98::mNeonGreen (Figure 7A). Line traces were drawn to pass through the central plane of nuclei identified by Hoechst staining. For each nucleus, the two peak values for the nuclear envelope rim were averaged and normalized to the average value of Nup62 in the nucleoplasm. To quantify the partitioning of TFEB::GFP, IBB_domain_::mNeonGreen, TDP-43::wrmScarlet, CRM1::mNeonGreen, and G3BP::mCherry between the nucleus and cytoplasm (Figures 5E and S7E) line traces were drawn to pass through the cytoplasm as well as the nucleoplasm and image background. Average intensity values for the nucleus and cytoplasm were background subtracted, then the value for the nucleus was divided by that of the cytoplasm.

To quantify cytoplasmic levels of mNeonGreen::Nup358 in oocytes versus somatic cells and embryos (Figure 1D), single focal planes captured from the same animal were exported to ImageJ. For each animal 3 ROIs in the -1 oocyte, somatic cell type of interest, or 4-cell embryo cytoplasm were measured, averaged, and normalized to the image background. Values for the somatic cell or embryo cytoplasm were then normalized to that of the oocyte within the same animal.

### Statistical analysis

All statistical tests were performed using GraphPad Prism 9.2.0 software. For comparison of three or more groups, significance was determined using a one-way ANOVA. For comparison of two groups, significance was determined using an unpaired t-test. In all figures error bars represent 95% confidence intervals (CI). For all figures, ns indicates not significant; *, P <0.05; **, P <0.01; ***, P <0.001; ****, P <0.0001.

## Acknowledgements

We thank all members of the Seydoux and Cochella Labs, Orna Cohen-Fix, and the Baltimore Worm Club for support and many helpful discussions. We thank Cristina Ayuso for assistance with *C. elegans* genome engineering, Madeline Cassani for generating strain JH3656, and the Fromme Lab for generously sharing yeast strains and plasmids. We thank the Chuang Lab for sharing the transportin::mNeonGreen strain and the Greenstein Lab for sharing the GFP::NDC1 strain. Several *C. elegans* strains were provided by the Caenorhabditis Genetics Center (CGC), which is supported by the National Institutes of Health Office of Research Infrastructure Programs (P40 OD010440). This work was funded by the Spanish State Research Agency (PID2019-105069GB-I00) and the National Institutes of Health (R37HD037047). L.T. is a postdoctoral fellow of the Life Sciences Research Foundation supported by the Howard Hughes Medical Institute. G.S. is an investigator of the Howard Hughes Medical Institute.

## Author contributions

L. Thomas and B. Taleb Ismail: investigation, formal analysis, validation, and visualization. L. Thomas and G. Seydoux: conceptualization and writing – original draft. P. Askjaer: writing – review and editing. G. Seydoux and P. Askjaer: funding acquisition and supervision.

## Supplemental videos

**Video S1. Nup foci, marked by GFP::Nup88, fully disassemble during mitosis.** Representative time-lapse images of CRISPR-tagged GFP::Nup88 (grey) with a mCherry::histone transgene (red) in a *C. elegans* embryo. Images are 15 μm maximum intensity projections at 2 min intervals.

**Video S2. Nup foci, marked by mNeonGreen::Nup98, fully disassemble during mitosis.** Representative time-lapse images of CRISPR-tagged mNeonGreen::Nup98 (grey) with a mCherry::histone transgene (red) in a *C. elegans* embryo. Images are 15 μm maximum intensity projections at 2 min intervals.

**Video S3. Swimming assay with control *C. elegans*.** Day 1 adults were placed in M9 media at room temperature and immediately imaged. This video is related to Figure 7C as well as Video S4 (Day 1 adults expressing *rab-3p*::Nup98::mNeonGreen). Video speed is real time.

**Video S4. Swimming assay with *C. elegans* expressing *rab-3p*::Nup98::mNeonGreen.** Day 1 adults were placed in M9 media at room temperature and immediately imaged. This video is related to Figure 7C as well as Video S3 (control Day 1 adults lacking *rab-3p*::Nup98::mNeonGreen). Video speed is real time.

---

**Table S1.**
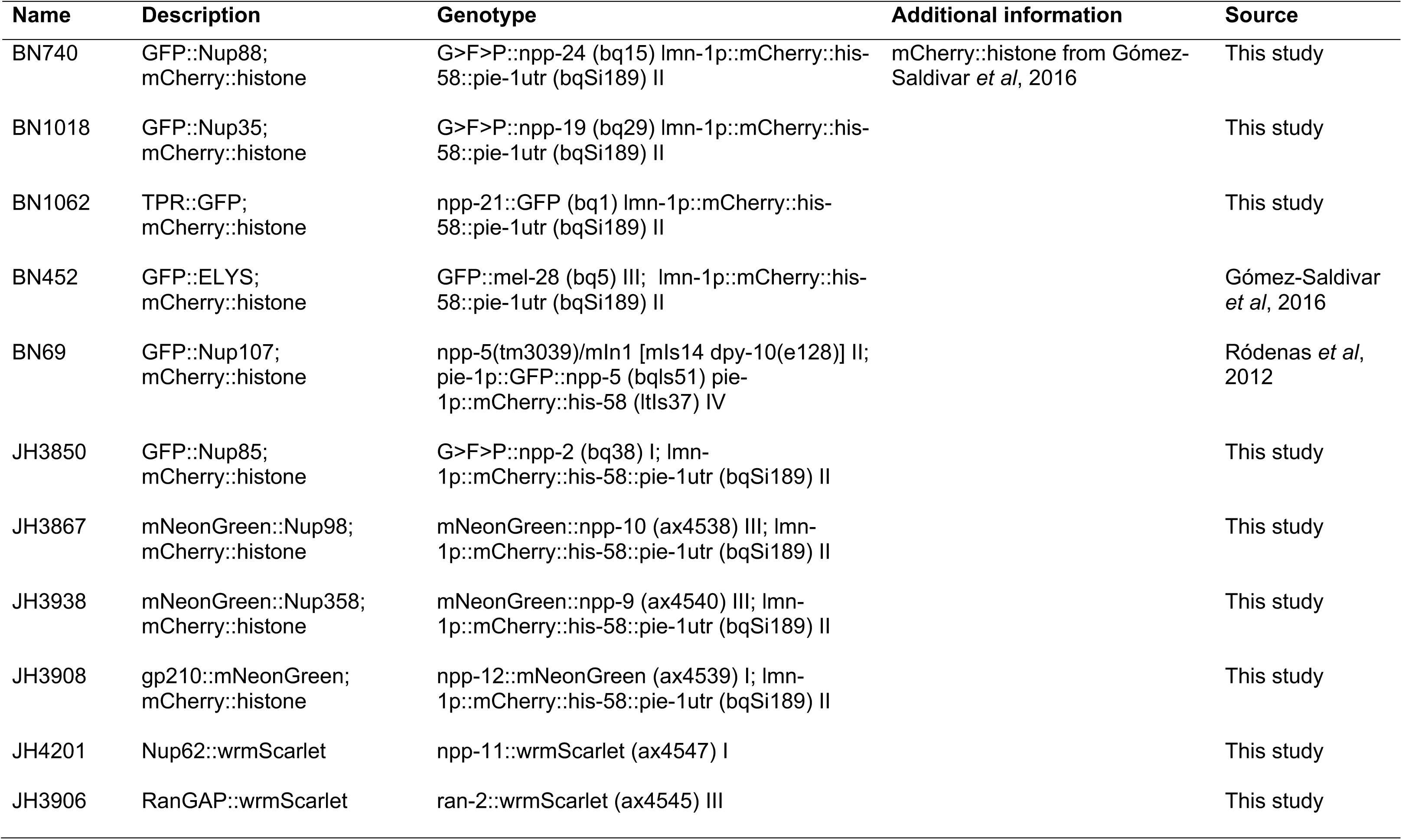
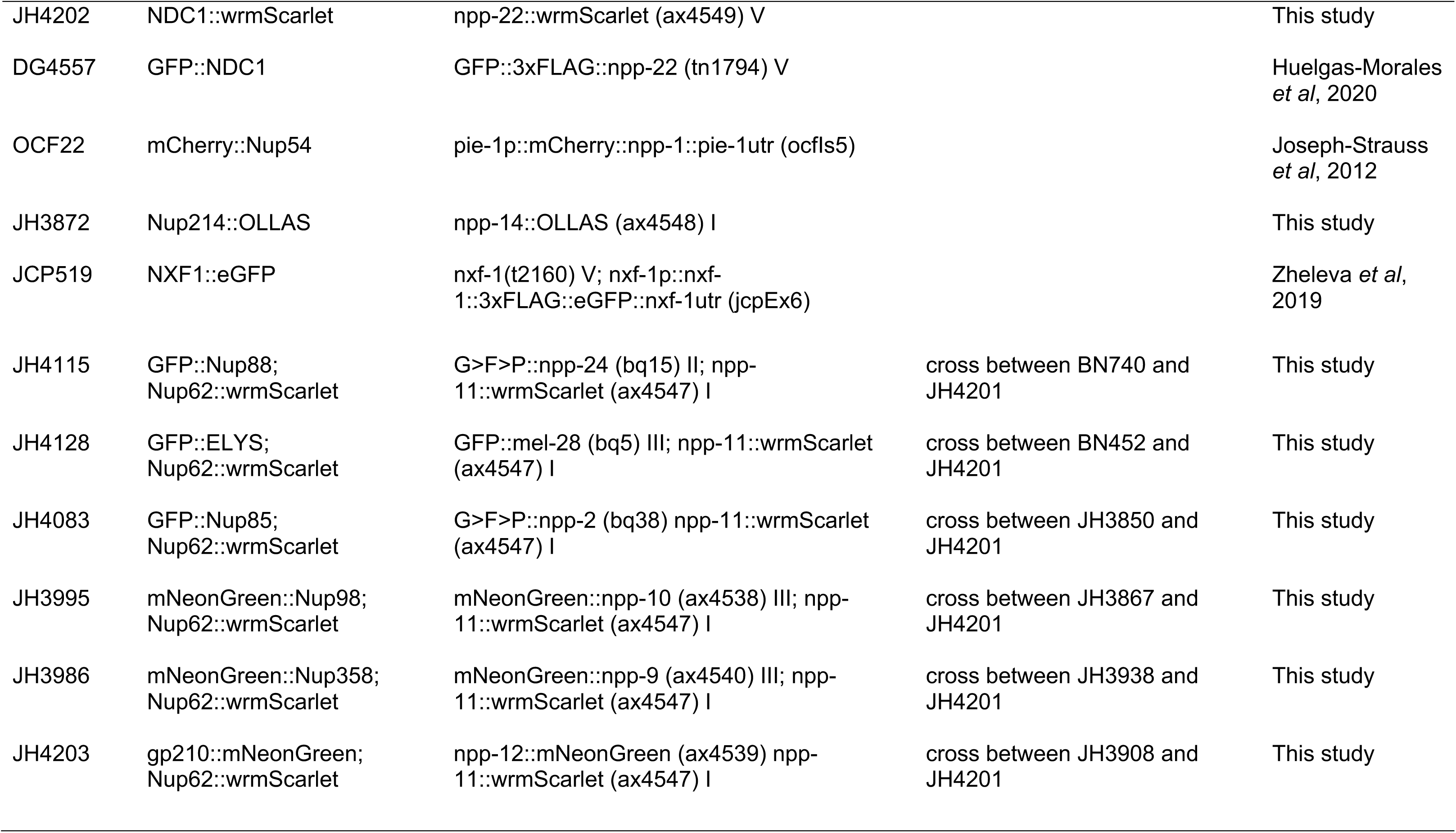
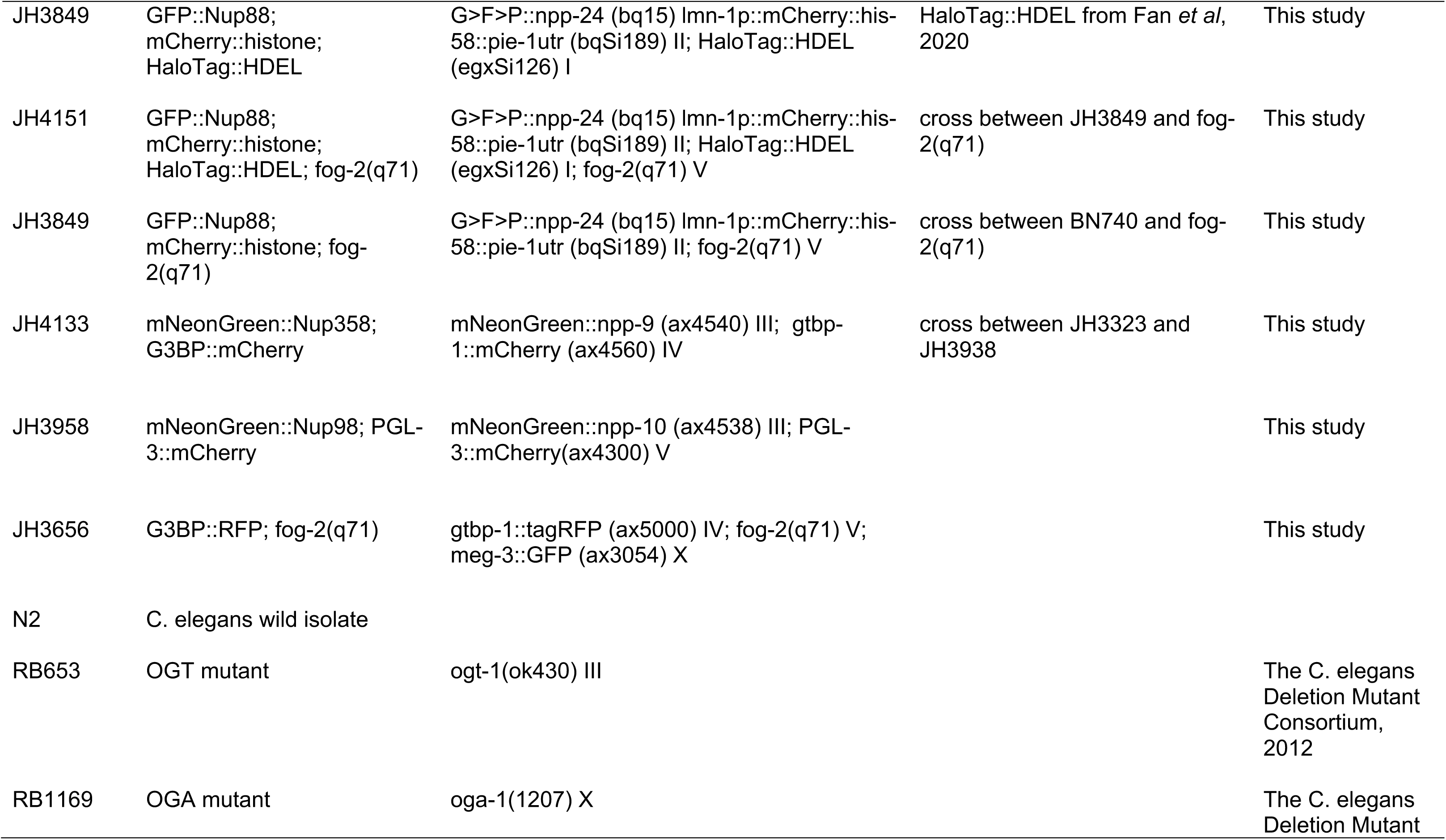
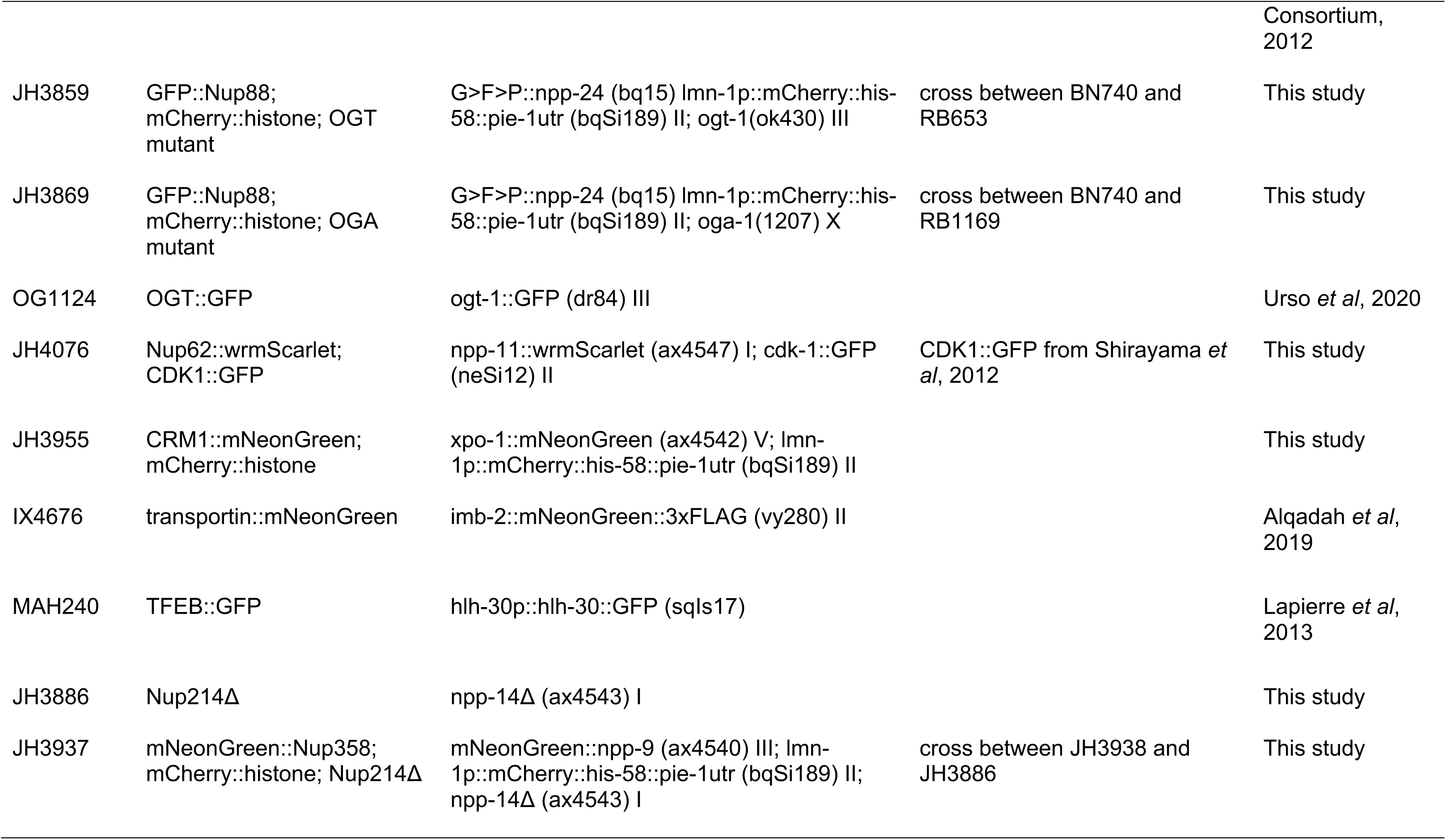
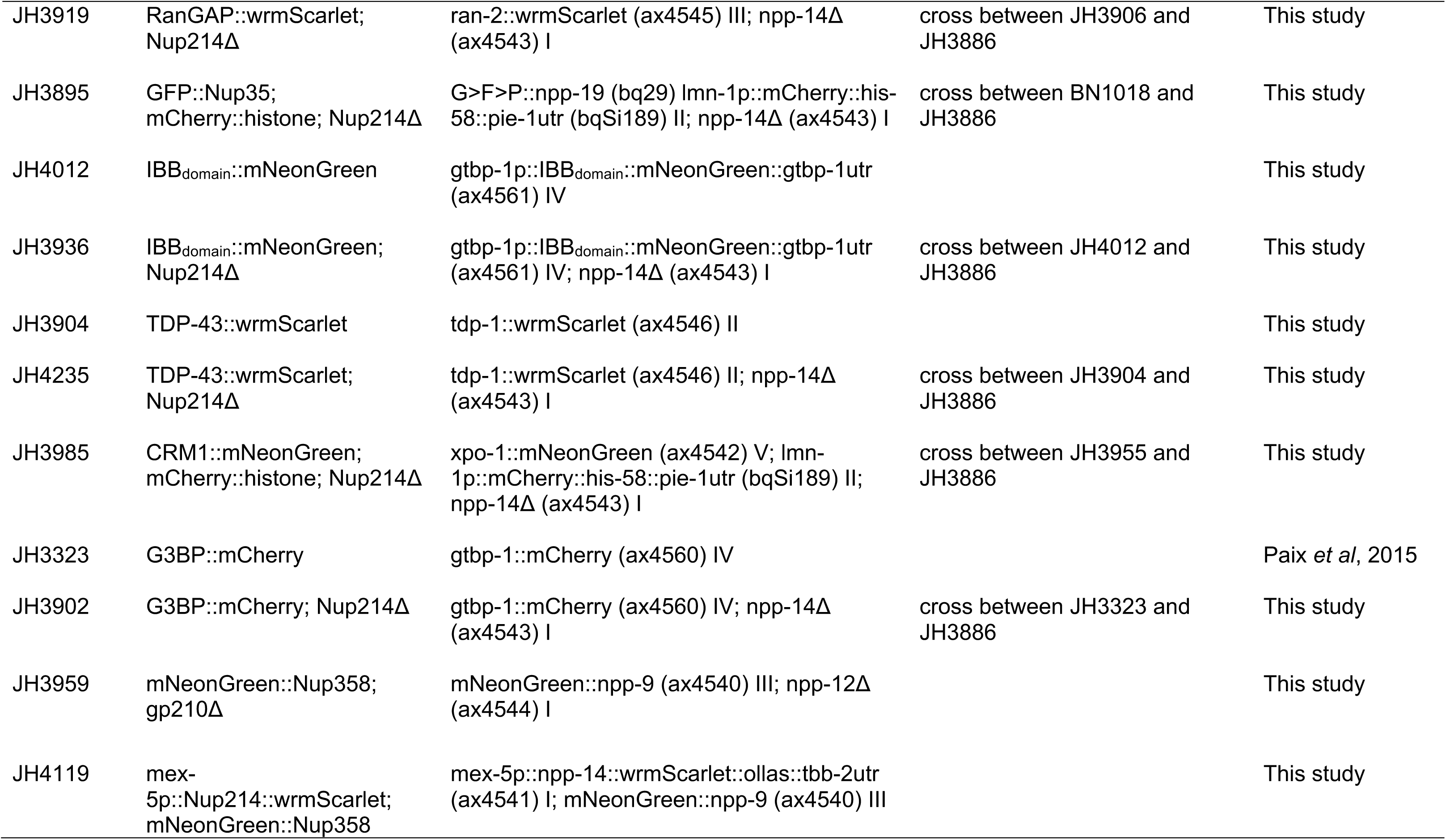
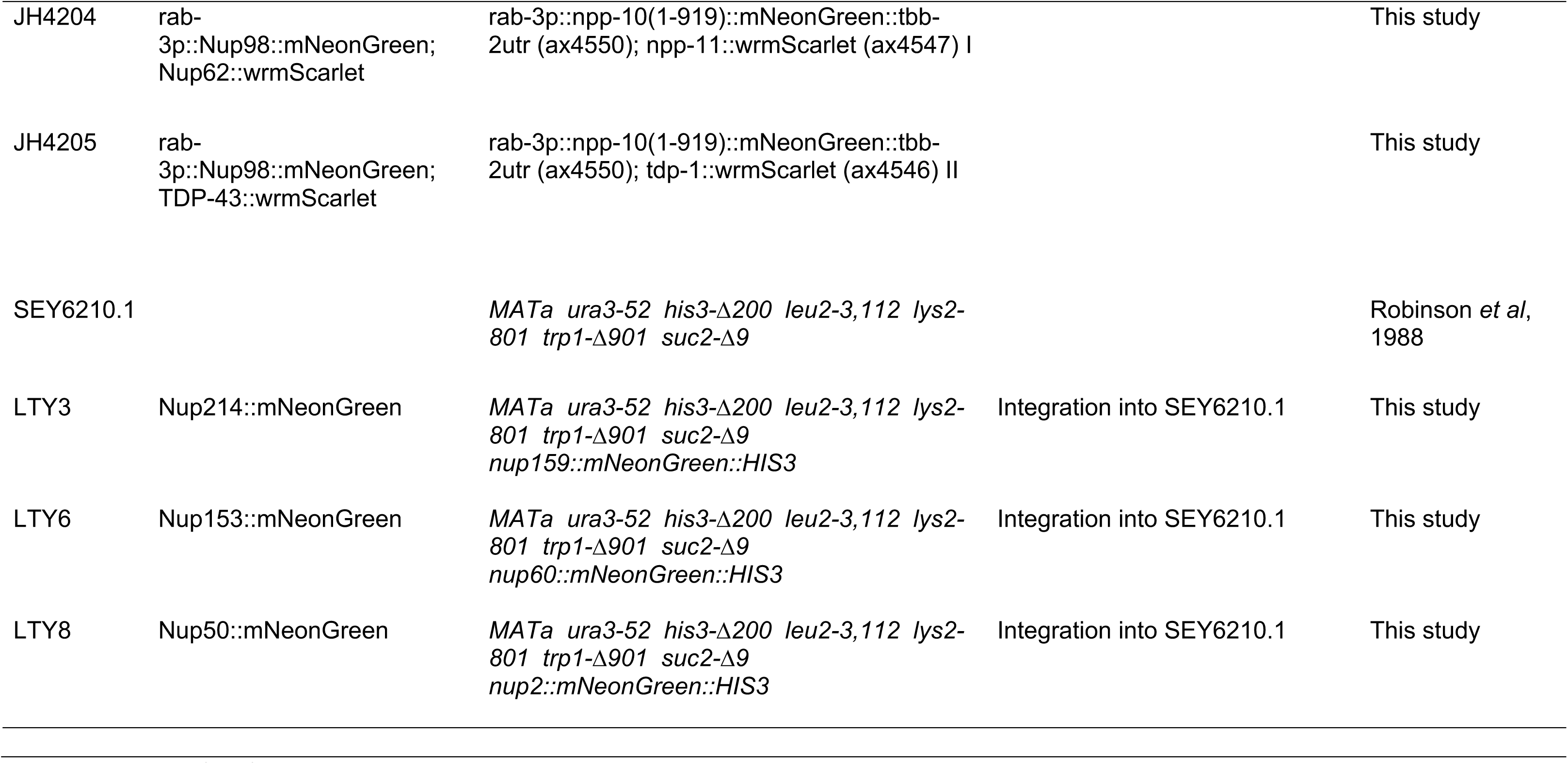
*C. elegans* and yeast strains used or generated in this study.

**Figure S1.**
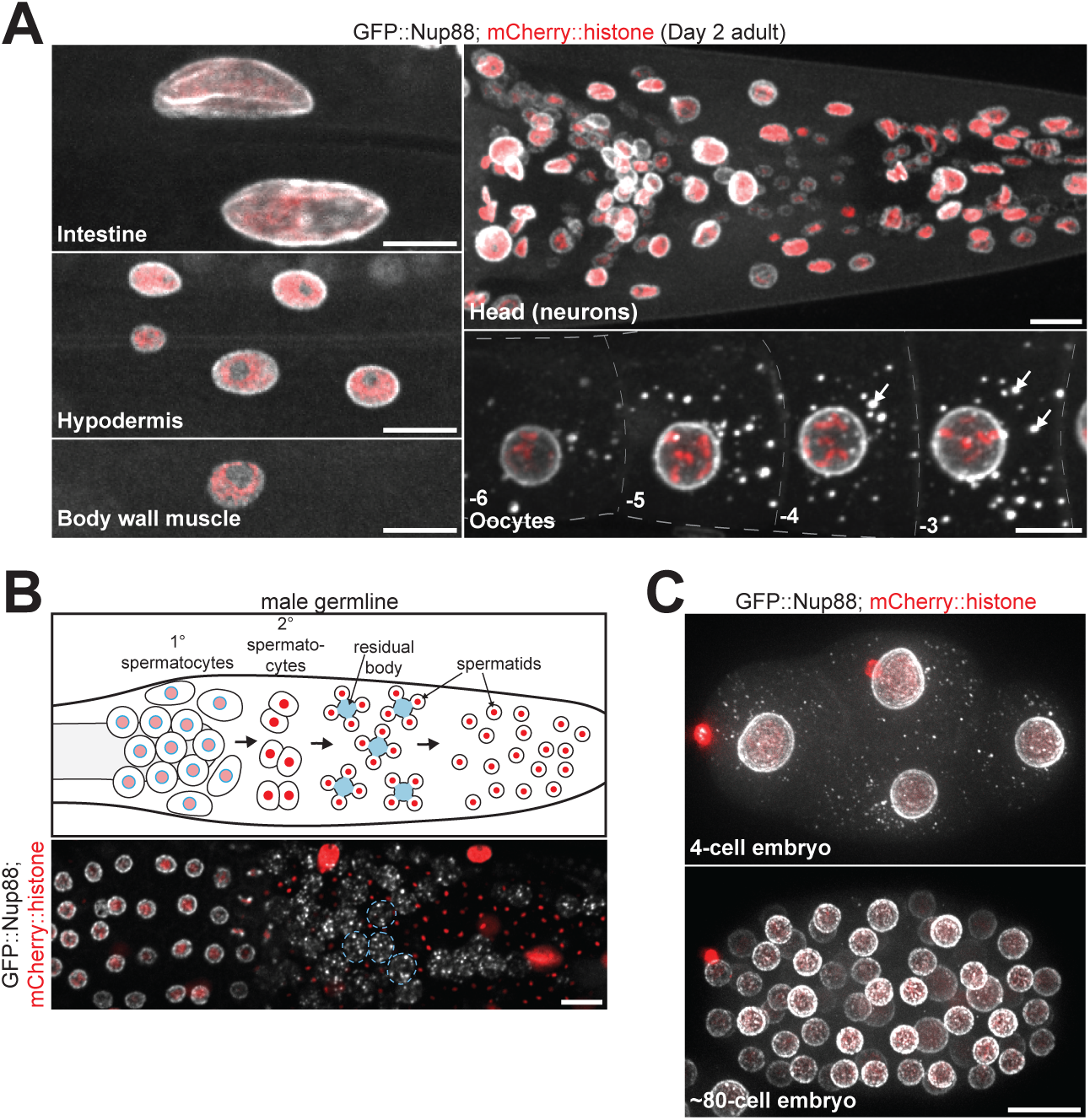
Cytoplasmic Nup foci are unique to germ cells and early embryos. A. Representative confocal micrographs of CRISPR-tagged GFP::Nup88 in the intestine, hypodermis, body wall muscle, head, and oocytes of Day 2 adult *C. elegans*. Nuclei are marked by a mCherry::histone transgene. White arrows denote cytoplasmic foci in oocytes. B. Top: Schematic depicting spermatogenesis in the *C. elegans* male germline. Following meiotic divisions spermatids bud from an anucleate residual body (denoted in blue); material within the residual body, including most Nups, is discarded. Bottom: Representative confocal micrograph showing GFP::Nup88 in the *C. elegans* male germline. Four residual bodies are outlined with a blue dashed line. C. Representative confocal micrographs showing GFP::Nup88 in interphase 4-cell versus ∼80-cell embryos. All images in this figure are maximum intensity projections. Scale bars = 10 μm.

**Figure S2.**
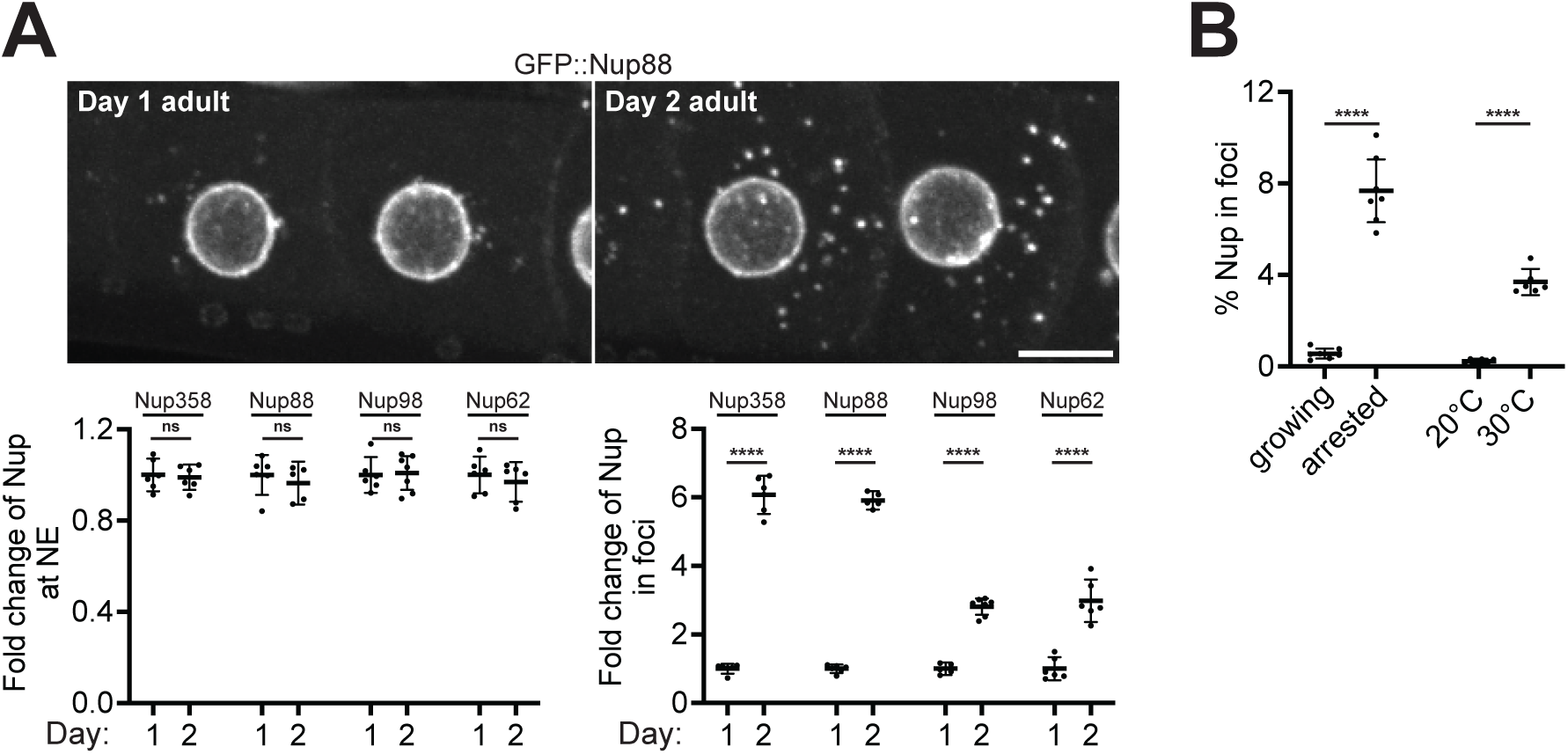
Nup foci formation increases significantly with age. A. Top: Representative confocal micrographs showing -3 and -4 oocytes of Day 1 versus Day 2 adults expressing CRISPR-tagged GFP::Nup88. Bottom left: Quantification of the total percent of each indicated Nup at the nuclear envelope (NE)/nucleoplasm in Day 1 versus Day 2 adults. Values are normalized so that the average value for Day 1 adults = 1.0. Error bars represent 95% CI for n > 5 germlines. Bottom right: Quantification of the total percent of each indicated Nup in foci in Day 1 versus Day 2 adults. Values are normalized so that the average value for Day 1 adults = 1.0. Error bars represent 95% CI for n > 5 germlines. Day 2 adult data is repeated from Figure 4A. B. Compiled quantification of the percent of Nup in foci in each indicated condition. Data correspond to micrographs in Figure 2A. Error bars represent 95% CI for n > 7 germlines (growing versus arrested) or n = 6 germlines (20°C versus 30°C). ****, P<0.0001; ns, not significant. All images in this figure are maximum intensity projections. Scale bar = 10 μm.

**Figure S3.**
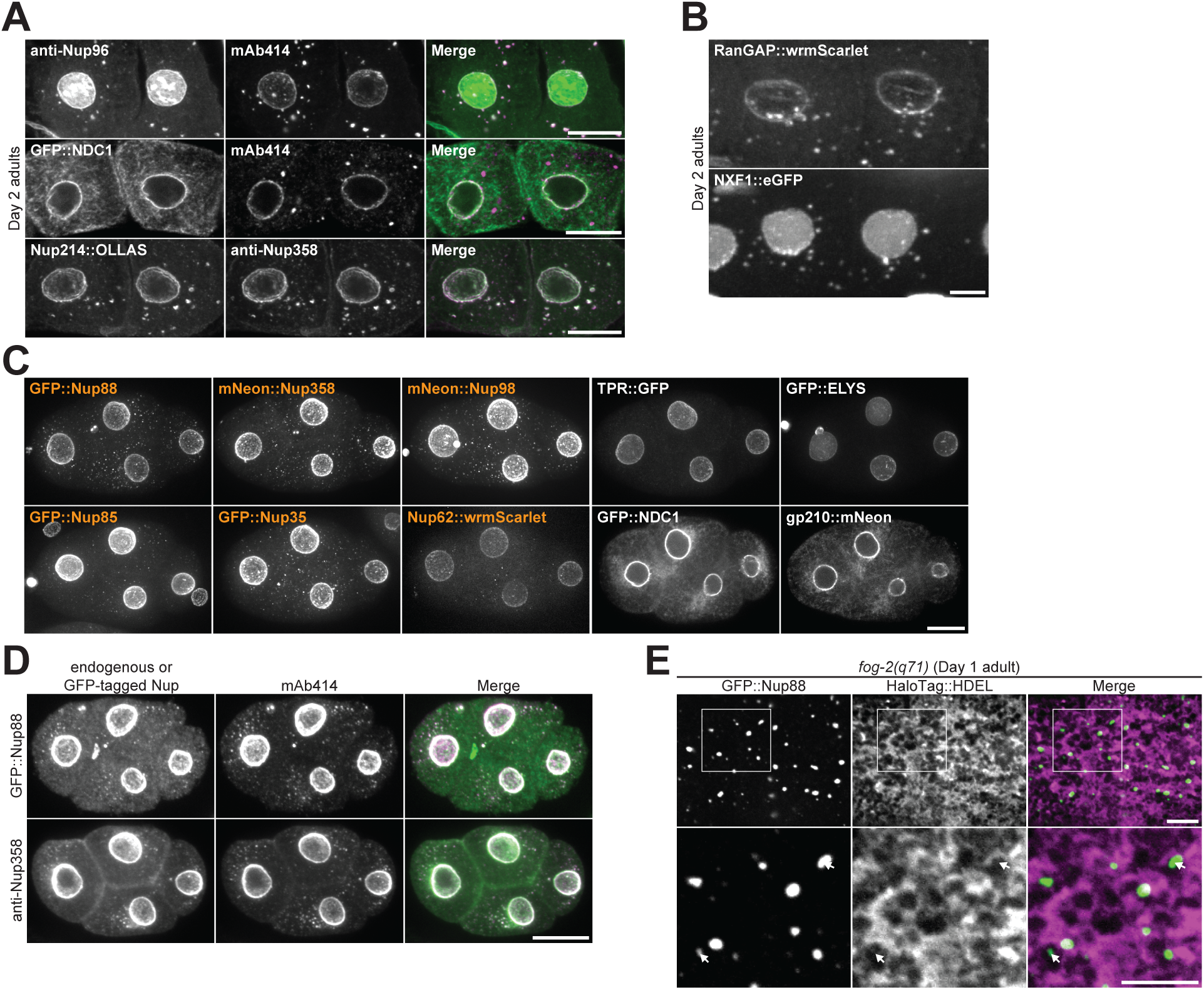
FG-Nups and their binding partners form cytoplasmic foci in early embryos. A. Top: Representative confocal micrographs depicting colocalization of endogenous Nup96 with mAb414 in Day 2 adult oocytes. Middle: Representative confocal micrographs depicting colocalization of CRISPR-tagged GFP::NDC1 with mAb414 in Day 2 adult oocytes. Bottom: Representative confocal micrographs showing colocalization of CRISPR-tagged Nup214::OLLAS with endogenous Nup358 in Day 2 adult oocytes. B. Representative confocal micrographs of -3 and -4 oocytes from Day 2 adults expressing CRISPR-tagged RanGAP::wrmScarlet or transgenic NXF1::eGFP. C. Representative confocal micrographs showing interphase 4-cell embryos with each designated CRISPR-tagged Nup. Orange labels denote Nups that localize to cytoplasmic foci. D. Representative confocal micrographs depicting colocalization of CRISPR-tagged GFP::Nup88 or endogenous Nup358 with mAb414 in 4-cell embryos. E. Representative confocal micrographs showing overlap of GFP::Nup88 with the luminal endoplasmic reticulum/nuclear envelope marker HaloTag::HDEL in a Day 1 adult *fog-2(q71)* arrested oocyte. 42% of foci completely overlapped with HaloTag::HDEL, 55% partially overlapped, and 3% showed no overlap with HaloTag::HDEL (n = 198). Areas indicated by white boxes are magnified below; white arrows indicate foci that do not completely overlap with the endoplasmic reticulum. All images in this figure are maximum intensity projections, with the exception of NDC1 with mAb414 (panel A), NDC1 and gp210 (panel C), and panel E which are single focal planes. Scale bars = 10 μm.

**Figure S4.**
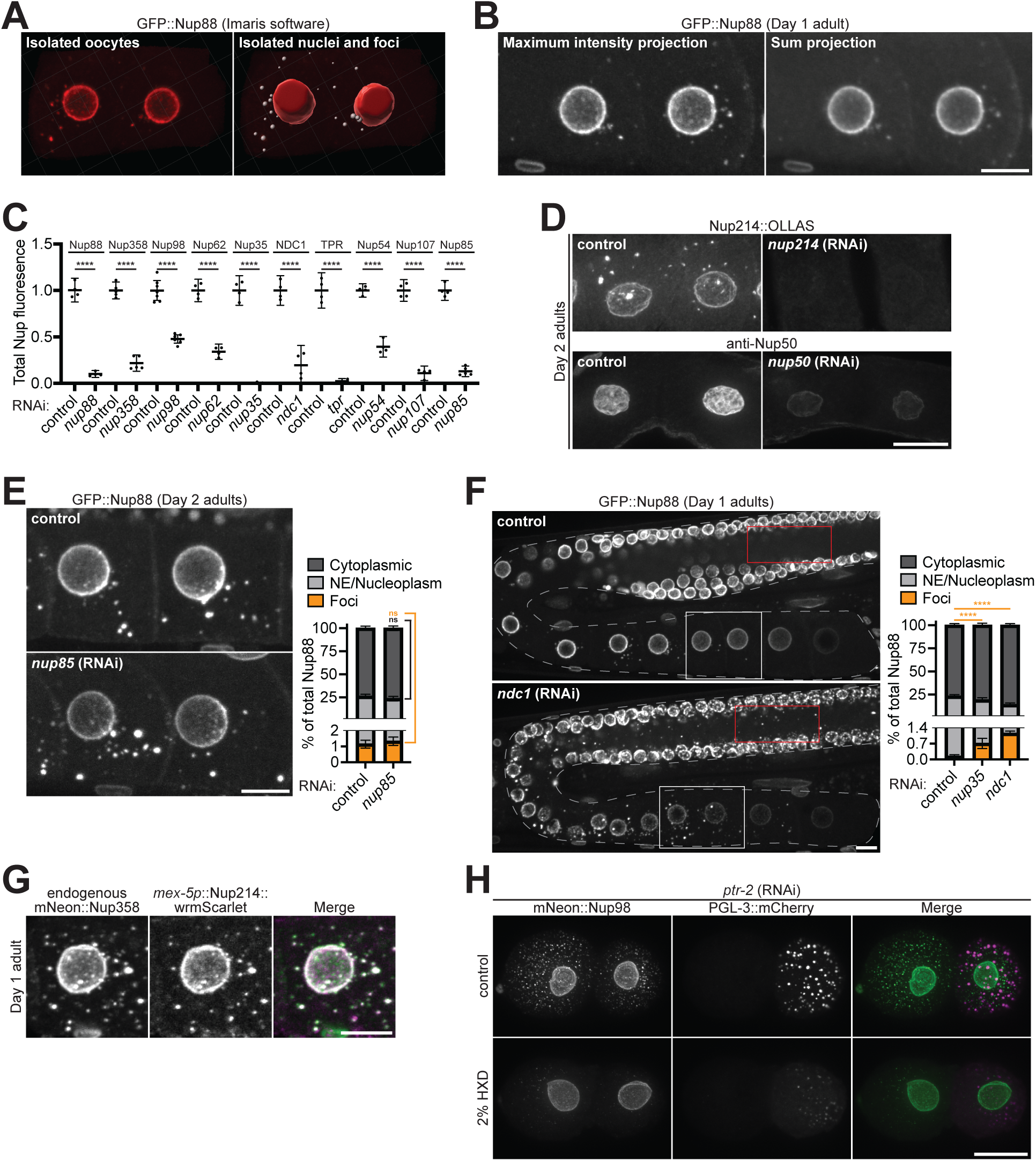
Nup foci are scaffolded by hydrophobic interactions. A. Images depicting the workflow used to measure the distribution of individual Nups in oocytes. Left: 3D reconstructions of -3 and -4 position oocytes were first isolated using Imaris software and total Nup fluorescence was measured. Right: Nuclei (red mask) and foci (grey mask) were then isolated, and the amount of Nup fluorescence for each group was measured. Also see materials and methods. B. Representative confocal micrographs showing CRISPR-tagged GFP::Nup88 in the -3 and -4 oocytes of a Day 1 adult. The 10 um Z stack is shown as a maximum intensity projection (left) or a sum projection (right). C. Quantification of the extent of RNAi-mediated depletion of each indicated Nup. Values are normalized so that the average control measurement = 1.0. Error bars represent 95% CI for n > 4 germlines. D. Representative confocal micrographs showing depletion of CRISPR-tagged Nup214::OLLAS or endogenous Nup50 following *nup214* or *nup50* RNAi, respectively. E. Left: Representative confocal micrographs showing -3 and -4 oocytes of Day 2 adults with GFP::Nup88 as a marker for foci formation, with control RNAi or following depletion of Nup85. Right: Quantification of the distribution of GFP::Nup88 between the cytoplasm (soluble), nuclear envelope (NE)/nucleoplasm, and cytoplasmic foci in control oocytes versus those depleted of Nup85. Error bars represent 95% CI for n = 6 germlines. F. Left: Representative confocal micrographs of Day 1 adult germlines expressing GFP::Nup88 with control RNAi or following depletion of NDC1. Red boxes denote the syncytial cytoplasm; foci are absent in control germlines but accumulate following depletion of NDC1. White boxes indicate the -3 and -4 oocytes used for quantification. Right: Quantification of the distribution of GFP::Nup88 between the cytoplasm, NE/nucleoplasm, and cytoplasmic foci in control oocytes versus those depleted of Nup35 or NDC1. Error bars represent 95% CI for n > 5 germlines. G. Representative confocal micrographs showing colocalization of CRISPR-tagged endogenous mNeonGreen::Nup358 with Nup214::wrmScarlet overexpressed using the *mex-5* promoter. H. Representative confocal micrographs showing CRISPR-tagged mNeonGreen::Nup98 and the P granule marker PGL-3::mCherry in 2-cell embryos treated or not with 2% 1,6-hexandiol (HXD) immediately prior to imaging. *ptr-2* RNAi was used to permeabilize the embryo eggshell to allow for HXD treatment. ****, P<0.0001; ns, not significant. All images in this figure are maximum intensity projections. Scale bars = 10 μm.

**Figure S5.**
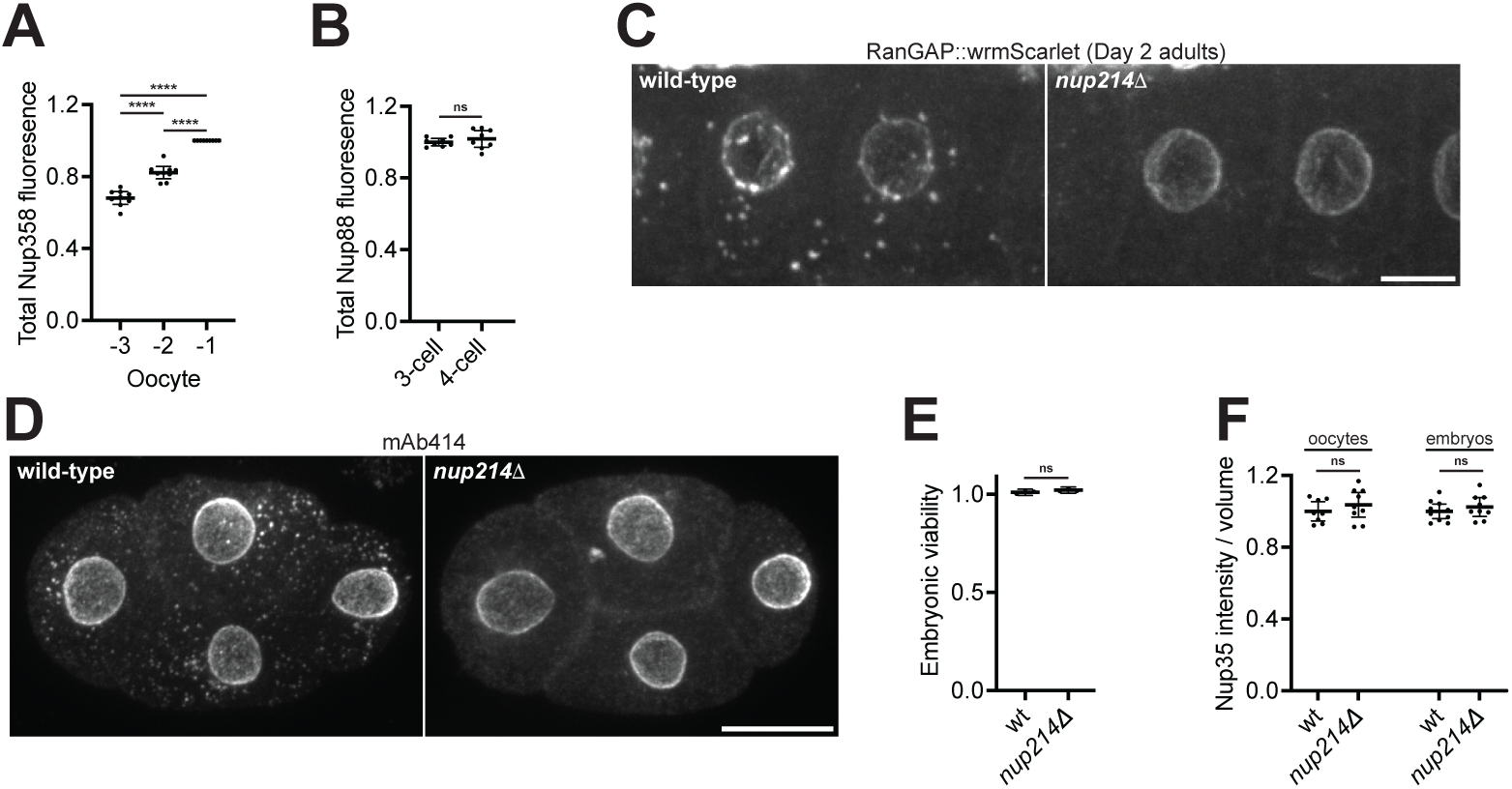
Robust Nup foci are not required for embryonic viability. A. Quantification of total mNeonGreen::Nup358 fluorescence in -3, -2, and -1 oocytes. Values are normalized within the same germline so that the value of the -1 oocyte = 1.0. Error bars represent 95% CI for n = 9 germlines. B. Quantification of total GFP::Nup88 fluorescence in 3-cell (mitosis) versus 4-cell (interphase) embryos. Values are normalized so that the average fluorescence of 3-cell embryos = 1.0. Error bars represent 95% CI for n > 6 embryos. C. Representative confocal micrographs showing CRISPR-tagged RanGAP::wrmScarlet in -3 and -4 oocytes of wild-type versus *nup214Δ* Day 2 adults. D. Representative confocal micrographs showing endogenous Nups, visualized by mAb414, in wild-type versus *nup214Δ* 4-cell embryos. E. Embryonic viability of wild-type *C. elegans* versus the *nup214Δ* mutant. Error bars represent 95% CI for N = 3 independent experiments with n = 907 (wild-type) or n = 892 (*nup214Δ*) animals. F. Quantification of the intensity of CRISPR-tagged GFP::Nup35 per nuclear volume in the -3 and -4 oocytes of Day 1 adults or 28-cell stage embryos. Values are normalized so that the average wild-type measurement = 1.0. Error bars represent 95% CI for n > 8 germlines and n > 9 embryos. ****, P<0.0001; ns, not significant. All images in this figure are maximum intensity projections. Scale bars = 10 μm.

**Figure S6.**
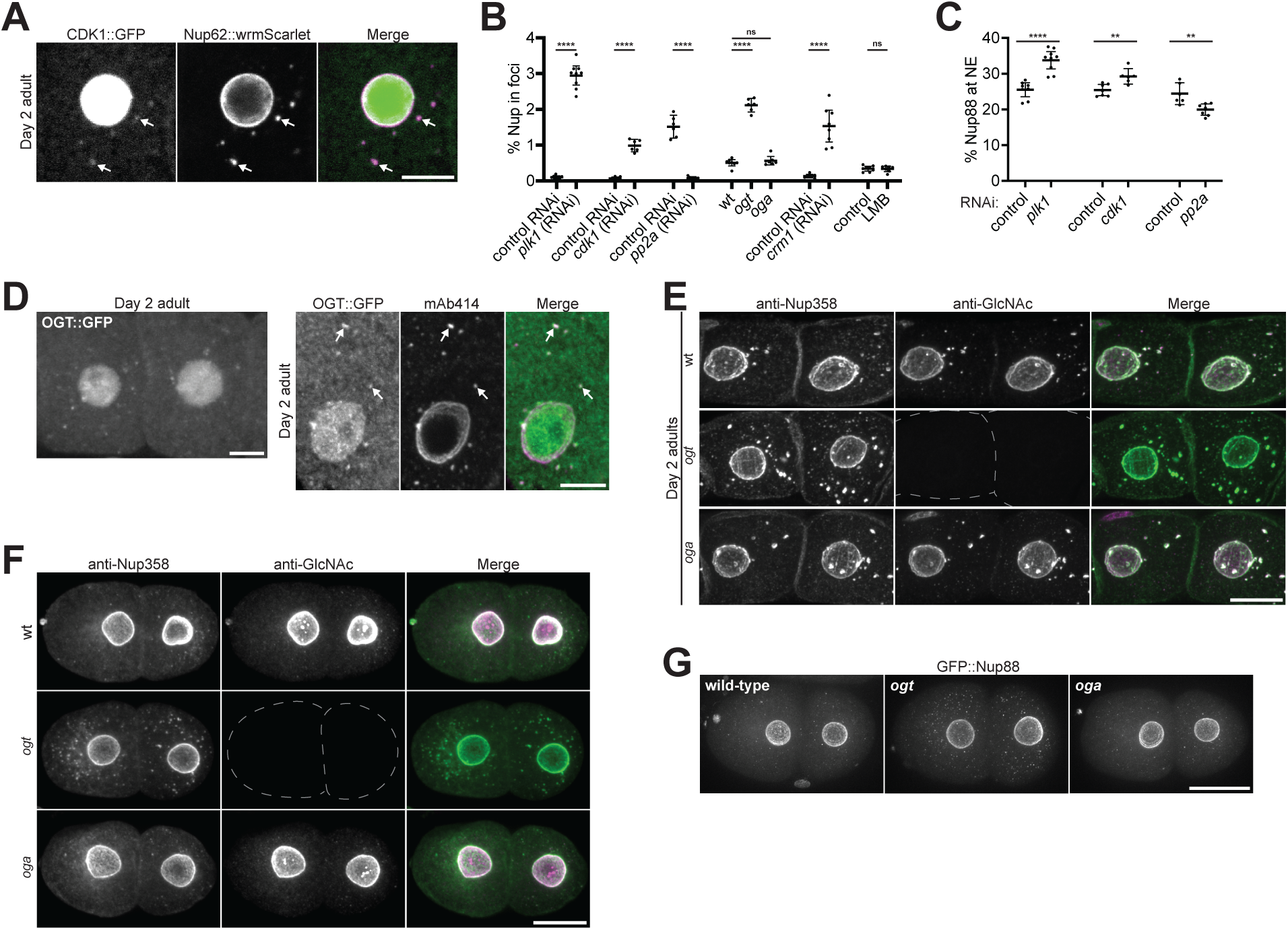
CDK1 and OGT localize to Nup foci and promote Nup solubility. A. Representative confocal micrographs showing colocalization of CRISPR-tagged Nup62::wrmScarlet with CDK1::GFP in a Day 2 adult oocyte. White arrows indicate overlap of CDK1 and Nup62 in cytoplasmic foci. B. Compiled quantification of the percent of Nup in foci in each indicated condition. Data correspond to micrographs in Figure 6B (*plk1* RNAi, n > 8 germlines; *cdk1* RNAi, n > 6 germlines; *pp2a* RNAi, n > 6 germlines), Figure 6C (*ogt* and *oga* mutants, n > 6 germlines), and Figure 6D (*crm1* RNAi, n > 7 germlines; LMB treatment, n > 8 germlines). C. Compiled quantification of the percent of GFP::Nup88 at the nuclear envelope (NE) under each indicated condition. Data correspond to micrographs in Figure 6B (*plk1* RNAi, n > 8 germlines; *cdk1* RNAi, n > 6 germlines; *pp2a* RNAi, n > 6 germlines). D. Left: Representative confocal micrograph showing CRISPR-tagged OGT::GFP in -3 and -4 oocytes of a Day 2 adult. Right: Colocalization of OGT::GFP with mAb414 in a Day 2 adult oocyte. White arrows indicate overlap of OGT and mAb414 in cytoplasmic foci. E. Representative confocal micrographs showing colocalization of endogenous Nup358 with the RL2 GlcNAc antibody in wild-type, *ogt*, or *oga* Day 2 adult oocytes. F. Representative confocal micrographs showing colocalization of endogenous Nup358 with the RL2 GlcNAc antibody in wild-type, *ogt*, or *oga* interphase 2-cell embryos. G. Representative confocal micrographs showing CRISPR-tagged GFP::Nup88 in wild-type, *ogt*, or *oga* 2-cell embryos. ****, P<0.0001; **, P<0.01; ns, not significant. All images in this figure are maximum intensity projections. Scale bars = 10 μm.

**Figure S7.**
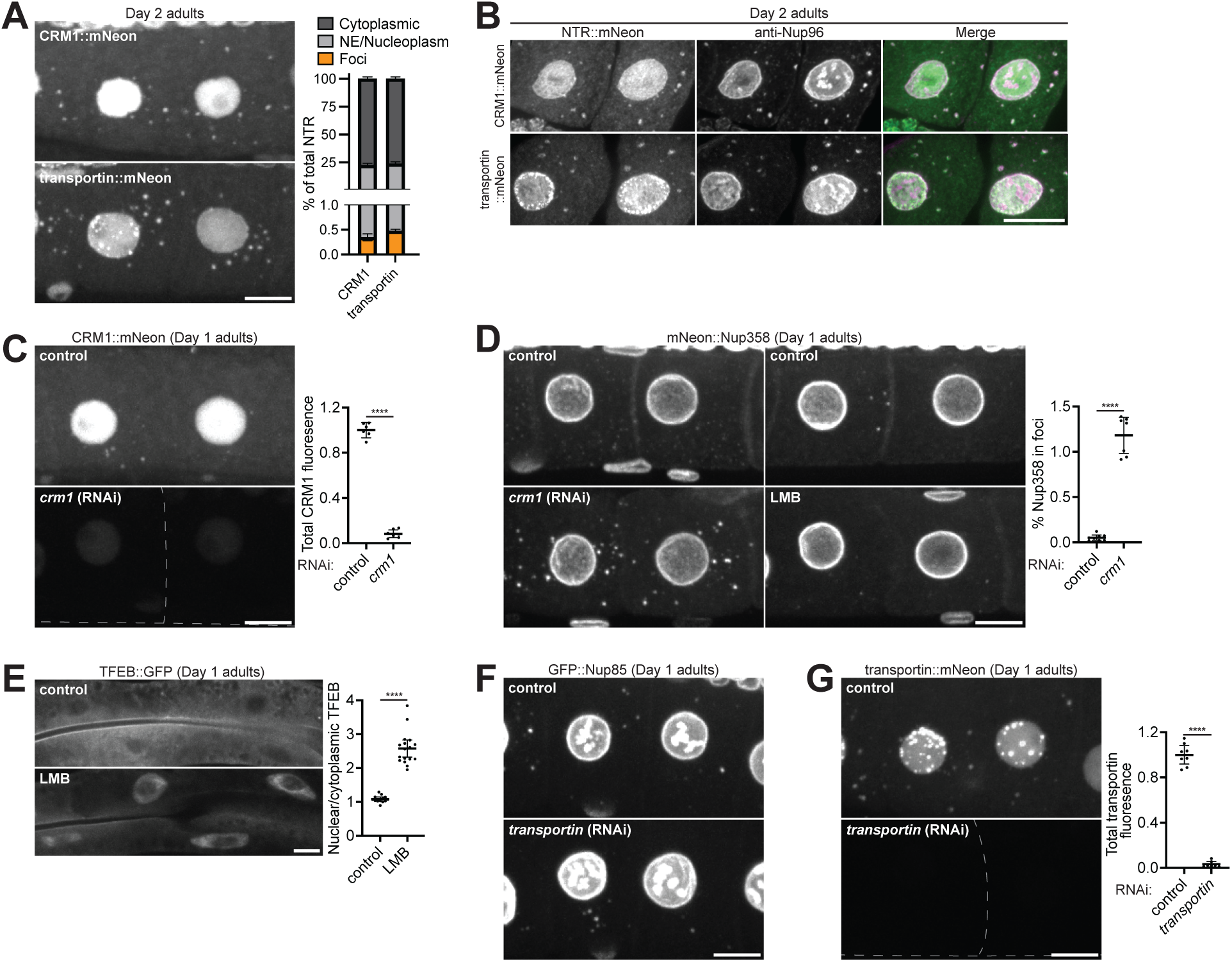
CRM1 and transportin localize to cytoplasmic Nup foci. A. Left: Representative confocal micrographs showing CRISPR-tagged CRM1::mNeonGreen or transportin::mNeonGreen in -3 and -4 oocytes of Day 2 adults. Right: Quantification of the distribution of CRM1::mNeonGreen and transportin::mNeonGreen between the cytoplasm (soluble), nuclear envelope (NE)/nucleoplasm, and cytoplasmic foci. Error bars represent 95% CI for n = 8 germlines. B. Representative confocal micrographs showing colocalization of CRM1::mNeonGreen and transportin::mNeonGreen with endogenous Nup96 in Day 2 adult oocytes. C. Left: Representative confocal micrographs showing CRM1::mNeonGreen in control -3 and -4 oocytes of Day 1 adults or oocytes targeted by *crm1* RNAi. Right: Quantification of total CRM1::mNeonGreen fluorescence in control oocytes or oocytes targeted by *crm1* RNAi. Values are normalized so that the average control measurement = 1.0. Error bars represent 95% CI for n > 6 germlines. D. Left: Representative confocal micrographs showing CRISPR-tagged mNeonGreen::Nup358 in control -3 and -4 oocytes of Day 1 adults or oocytes depleted of CRM1. Middle: Representative confocal micrographs showing mNeonGreen::Nup358 in control oocytes or following treatment with the CRM1 inhibitor leptomycin B (LMB). Right: Quantification of the total percent of mNeonGreen::Nup358 in foci in control oocytes or following CRM1 depletion. Error bars represent 95% CI for n > 7 germlines. E. Left: Representative confocal micrographs showing TFEB::GFP in control Day 1 adults or following LMB treatment. Right: Quantification of the nuclear/cytoplasmic ratio of TFEB::GFP in control cells or following LMB treatment; note that nuclear export of TFEB is mediated by CRM1 (Silvestrini *et al*, 2018). Error bars represent 95% CI for n > 13 nuclei. F. Representative confocal micrographs showing CRISPR-tagged GFP::Nup85 in -3 and -4 oocytes of control Day 1 adults or oocytes depleted of transportin. G. Left: Representative confocal micrographs showing transportin::mNeonGreen in control -3 and -4 oocytes of Day 1 adults or oocytes targeted by *transportin* RNAi. Right: Quantification of total transportin::mNeonGreen fluorescence in control oocytes or oocytes targeted by *transportin* RNAi. Values are normalized so that the average control measurement = 1.0. Error bars represent 95% CI for n > 7 germlines. ****, P<0.0001. All images in this figure are maximum intensity projections, with the exception of panel E which are single imaging planes. Scale bars = 10 μm.

**Figure S8.**
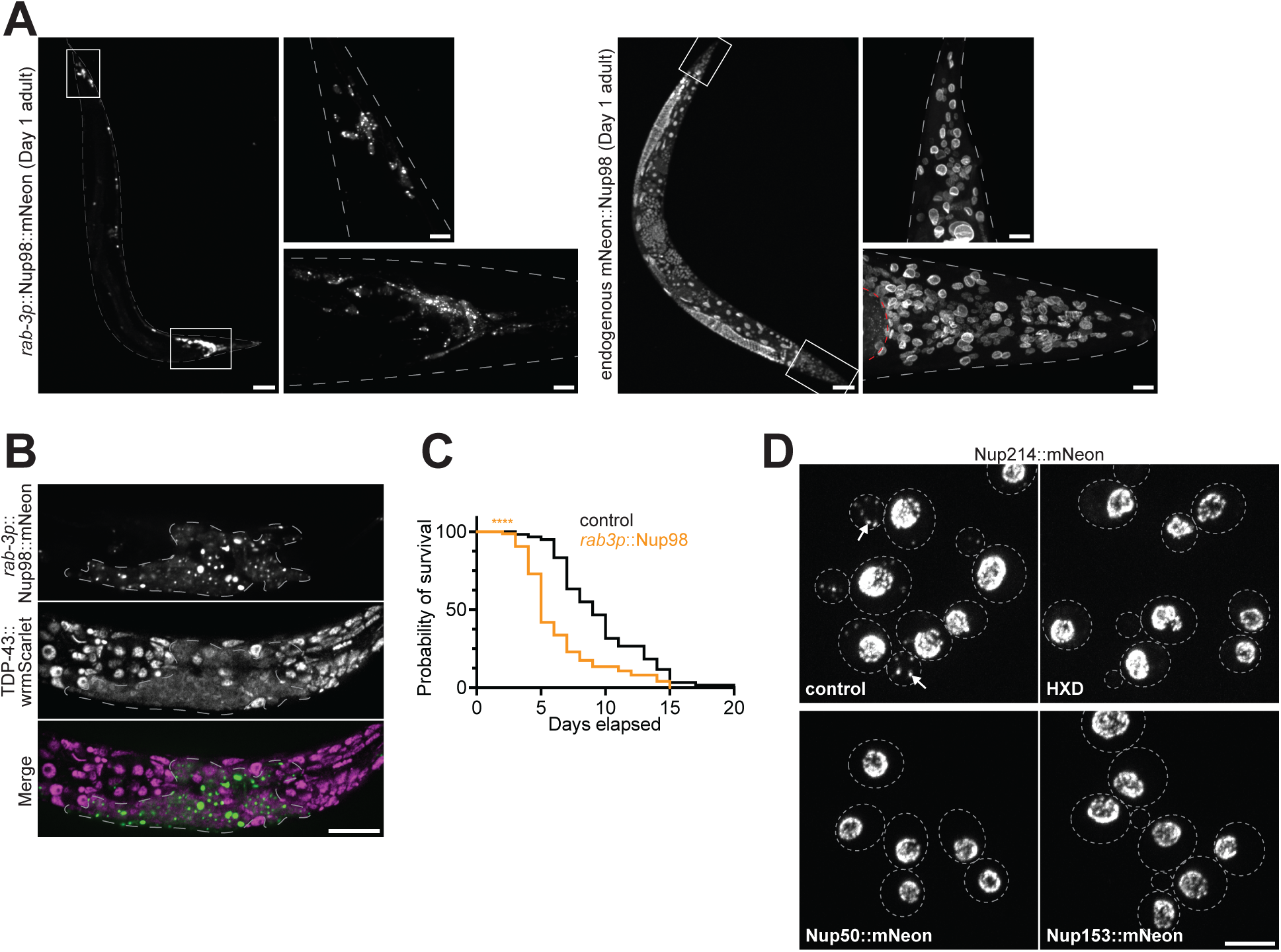
Ectopic Nup98 condensation in neurons is toxic. A. Left: Representative confocal micrographs showing transgenic *rab-3p*::Nup98::mNeonGreen in a Day 1 adult *C. elegans*; image was acquired using 30% laser power. The head and tail ganglia indicated by white boxes are magnified in the panels to the right. Right: Representative confocal micrographs showing endogenous CRISPR-tagged mNeonGreen::Nup98 in a Day 1 adult *C. elegans*; image was acquired using 100% laser power. The head and tail indicated by white boxes are magnified in the panels to the right. Red dashes outline part of the germline. B. Representative confocal micrographs showing localization of CRISPR-tagged TDP-43::wrmScarlet and transgenic *rab-3p*::Nup98::mNeonGreen in the head of an L1 larva. Note that transgenic Nup98::mNeonGreen is only expressed in neurons; dashed line indicates cells with Nup98::mNeonGreen expression. C. Survival curve of control *C. elegans* versus those with ectopically expressed *rab-3p*::Nup98::mNeonGreen. n = 61 (control) or n = 74 (*rab-3p*::Nup98::mNeonGreen) animals. D. Representative confocal micrographs showing endogenous Nup214, Nup50, and Nup153 tagged with mNeonGreen in yeast cells. Yeast expressing Nup214::mNeonGreen were treated with 5% 1,6-hexandiol (HXD) for 10 min prior to imaging. White arrows denote cytoplasmic Nup foci. ****, P<0.0001. All images in this figure are maximum intensity projections. Scale bars = 100 μm (panel A, whole worms), 5 μm (panel D), or 10 μm (all other panels).

